# Autoimmunity and *H. pylori* infection cooperatively promote epithelial tumorigenic differentiation and systemic immune modulation

**DOI:** 10.1101/2025.11.23.690054

**Authors:** Emily N. Vazquez, Clarisel Lozano, Amanda M. Honan, Elena Zaika, JeanMarie Houghton, Wael El-Rifai, Alexander Zaika, Zhibin Chen

## Abstract

**Background and Aims:** Gastric cancer (GC) demographics have shifted, with a growing impact on younger populations, particularly women. Chronic *Helicobacter pylori* (HP) infection is the leading risk factor for GC; however, decreasing HP prevalence means it is unlikely the cause of rising incidence of early-onset GC. Autoimmunity, which is more prevalent in women, has been proposed as a contributing factor. As autoimmunity prevalence increases, HP and gastric autoimmunity are likely to converge within one individual. This study aimed to examine how combinatorial HP infection and autoimmunity affect tumorigenesis and the immune landscape.

**Methods:** A mouse model mimicking human CTLA4 insufficiency and its risk of autoimmunity-driven GC development, CTLA4KD, was infected with HP strain PMSS1 by oral gavage. Tumorigenesis was histologically assed. Hematopoietic and non-hematopoietic cells were analyzed using 40-color spectral flow cytometry. Human PBMCs were co-cultured with HP, followed by flow cytometry and qRT-PCR for gene expression of sorted PBMC subsets.

**Results:** HP infection in CTLA4KD mice advanced dysplasia and increased tertiary lymphoid structures within the gastric mucosa. Loss of E-Cadherin and DMBT1 was exacerbated in the combinatorial setting of autoimmunity and HP infection. In the innate immune system HP and autoimmunity showed an additive effect in systemic NK cell decreases. In the adaptive immune system, HP and autoimmunity cooperatively elevated the proportion of CD4^+^Foxp3^-^ cells expressing the transcription factor Helios, persisting 6-8 months post infection. The Helios-expressing CD4 cells co-expressed inhibitory markers PD-1, CD200 and CD39.

**Conclusion:** Autoimmunity and HP infection cooperatively elicit long-lasting suppression of innate and adaptive immunity while promoting epithelial tumorigenic differentiation in the stomach.

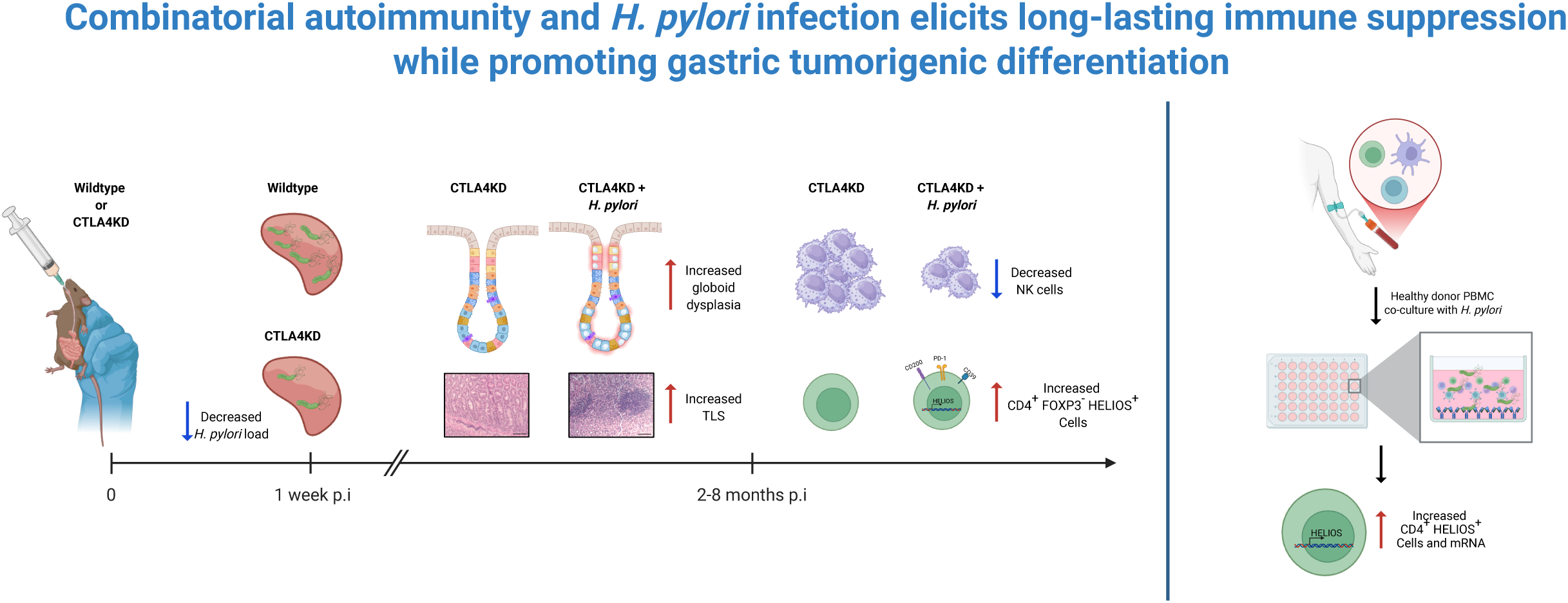

**Background and context:** Autoimmunity is emerging as a contributing factor for early-onset gastric cancer especially in young women, but it is unknown how it interacts with the most prevalent etiology factor, *Helicobacter pylori*.

**New findings:** This study provides the initial evidence that *Helicobacter pylori* and autoimmunity cooperatively cause long-lasting suppression of innate and adaptive immunity while promoting epithelial tumorigenic differentiation in the stomach.

**Limitations:** Even with the murine-adapted *Helicobacter pylori* strain PMSS1, a human pathogen, the mouse modeling remains technically and conceptually challenging and can only capture some aspects of human *Helicobacter pylori* infection.

**Clinical research relevance:** This study showcases how autoimmunity predisposed by host genetics may cooperate with a prevalent environment factor, *Helicobacter pylori,* to promote gastric cancer development, with cellular and histopathological changes in the stomach, as well as local and systemic alteration of immune profiles. The lasting effects long after the elimination of *Helicobacter pylori* suggest the necessity of remedies besides *Helicobacter pylori* eradication.

**Basic research relevance:** This study provides molecular and cellular clues for how a convergence of autoimmunity and *Helicobacter pylori* cooperatively downregulates tumor suppressors in gastric epithelial cells while causing long-lasting suppression of innate and adaptive immunity. More studies are needed for better understanding of the interaction between *Helicobacter pylori* and autoimmunity, to help develop interventions against the alarming trend of early-onset gastric cancer.

**Lay summary:** This study provides initial insight into how autoimmunity and *Helicobacter pylori* in combination may impinge on multi-pronged effect on the stomach and the immune system to accelerate gastric cancer development.

## Introduction

GC is the fifth most common and fourth most deadly cancer worldwide^1^. While many factors contribute to GC, the main risk factor for non-cardia GC is *Helicobacter pylori* (HP) infection^2^, a type I carcinogen, as characterized by the World Health Organization^3^.

Despite declining HP infection rates, GC incidence is not dropping proportionally, suggesting a rise in HP negative GCs^4^. Moreover, cancer incidence is shifting towards younger individuals, particularly those under 50^5^. While environmental factors, such as smoking and diet^6^, may contribute to the surge in younger individuals, autoimmunity has been implicated in increased non-HP GC^7^, especially for the faster rising cancer rates in women, a group with higher risk of autoimmune disease^8^. Thus, autoimmunity-initiated GC warrants careful examination as an etiological factor for early-onset GC.

Autoimmunity results from the breakdown of immune tolerance. Among its mechanisms, peripheral regulation by immune checkpoints, such as CTLA4, plays a key role^9^. CTLA4 haploinsufficient patients present with autoimmune disorders and elevated cancer risk^10^, including GC^11, 12^. To model CTLA4 haploinsufficiency, our group developed a transgenic CTLA4 shRNA (CTLA4KD) mouse that reduces CTLA4 expression by ∼60%. CTLA4KD mice develop autoimmune gastritis and spasmolytic polypeptide-expressing metaplasia (SPEM) by 5-6 weeks of age which progresses spontaneously to GC after 12 months of age^13^.

While host factors are increasingly scrutinized, HP remains the leading cause of GC. HP carcinogenicity is largely driven by virulence factors, such as CagA that is transformative to cells^3^, and other virulence factors that elicit chronic inflammation^14^.

As autoimmune disease incidence rises, it is probable that the effects of HP infection and autoimmunity converge on the gastric epithelium, with similar or distinct impact. A recent study has shown that autoimmune gastritis and HP can cause similar types of metaplasia^15^ However, it remains unclear how the distinct inflammatory conditions of HP and autoimmunity interact to affect tumorigenesis. In this study, we investigated how the combination of HP infection and gastric autoimmunity impacts the immune microenvironment and the initiation of tumorigenesis.

## Materials and Methods

### Mice

Male and female mice were used as no sex-differences were observed. Mice were infected with 1×10^9^ colony-forming units (CFU) of HP strain PMSS1 in 500μl of Brucella broth by oral gavage for two consecutive days. Mice may be fasted for up to 6 hours prior to infection. Control mice given Brucella broth by oral gavage for two consecutive days were combined with untreated mice, as no difference was detected. CTLA4KD mice were previously described^13^. Mice used in the study were on either a C57BL/6 (B6) or Balb/c x C57BL/6 F1 (CB6F1) genetic background. The studies were approved by the Institutional Animal Care and Use Committee at the University of Miami.

### Quantification of H. pylori load

One week post HP infection stool was collected from the mice. DNA was isolated using the QIAamp Fast DNA Stool Mini Kit (Qiagen) according to manufacturer’s protocol. HP was quantified by quantitative RT-PCR using primers and a probe specific for the HP Urease B gene: 5’- AGATGGCAAAATCGCTGGCA-3’ (forward), 5’- CGGCTAAGGCTTCAGTAGCAG-3’ and 5’-TGGTAAAGGCGGTAACAAAGACATGCA-3’ (probe), adapted from previously described primer and probe^16^. Results were compared to a standard for quantification.

### Culture and Urease test for gastric HP detection at sacrifice

After euthanasia, mouse stomachs were homogenized in PBS containing nalidixic acid, vancomycin, amphotericin, bacitracin and polymyxin B. Homogenized samples were plated on trypticase soy agar plates supplemented with 5% sheep’s blood plates and incubated at 37° C with 5% CO_2_ for one week. Following incubation, urease test media was inoculated with swabbed samples from the plates and incubated overnight at 37°C with 5% CO_2._ Changing of the urease media color from yellow to pink indicates a positive result.

### Stomach processing into single-cell preparations

Mouse gastric mucosa was collected after euthanasia, cut into pieces <5mm in size and incubated in RPMI media supplemented with FBS and pen/step, containing Collagenase IV (1mg/ml) and DNase (10 U/ml) for 20 minutes at 37° C on a shaker. Following incubation supernatant was collected, and more media was added and subsequently incubated for 20 minutes at 37° C. This was repeated twice for a total of three 20-minute incubations. Digested tissue was further mechanically dissociated using an 18G needle and filtered using a 70μm filter and washed with PBS.

### Full spectrum flow cytometry

A 40-color flow cytometry panel was designed to comprehensively characterize the hematopoietic and non-hematopoietic compartments. Single-cell preparations from the gastric lymph nodes (GLN), mesenteric lymph nodes (MLN), spleen (SPL) and stomach mucosa (STO) underwent flow cytometry. Cells were blocked against nonspecific antibody binding using normal mouse serum, anti-CD16/32 antibody, BD Horizon Brilliant Stain Buffer Plus and CellBlox blocking buffer. Following blocking, cells were first stained with antibodies for cell surface markers, and then fixed and permeabilized for intracellular staining using Transcription Factor Buffer Set (BD Biosciences. Antibodies used are listed in supplemental table 1. Dead cells were stained and excluded using Zombie NIR staining. Doublets were gated out from analysis. Manual gating was conducted using FCS express software.

### viSNE and SPADE clustering analysis

Spectral flow cytometry data underwent viSNE dimensionality reducing analysis on the Cytobank platform using FCS files were pre-gated on live cells (Zombie NIR^-^) and a hematopoietic lineage (CD45) cells using FCS express software. viSNE analysis transformed FCS data into 2 dimensions using the Barnes-Hut implementation of the t-distributed stochastic neighbor embedding (tSNE) algorithm. Analysis was performed on the samples using proportional sampling, with 7,500 iterations, a perplexity of 30, and a θ of 0.5, following Cytobank user guides and similar procedures reported in a previous study^17^. The following markers were used as clustering channels: CD11b. CD11c, CD19, CD4, CD8a, Foxp3, Gata3, IgD, MHCII, NK1.1, RORγt, T-bet, TCRβ. Following viSNE analysis Spanning-tree Progression Analysis of Density-normalized Events (SPADE)^18^ clustering analysis was conducted on the results. Cells were clustered on the dimensionality reduction channels tSNE1 and tSNE2. SPADE trees were manually analyzed, and nodes were gated based on marker expression. Gates were then overlaid on the viSNE plots for visualization.

### In vitro human PBMC and HP co-culture

24-well suspension plates were coated with anti-CD3 and anti-CD28 antibodies in PBS for T cell stimulation. Two million human PBMCs were added to each well following coating of the plate in antibiotic free RPMI supplemented with 10% FBS and L-glutamine. De-identified healthy PBMCs were obtained from a blood bank located in the same geographic area for which the university hospital serves, and the donors were screened for health status and tested for infectious agents per the standard blood bank policy. Three days after the cells were stimulated, HP (PMSS1) at a MOI of 2 was added to cells and heat killed HP was used as a control. At 16 hours post infection cells were collected and analyzed by flow cytometry.

### Human PBMC sorting and Helios quantitative RT-PCR

Following *in vitro* human PBMC and HP co-culture, samples were prepared for cell sorting. Cells were blocked for nonspecific antibody binding using normal mouse serum, anti-CD16/32 antibody in PBS with 2% FBS. Cells were then stained with fluorescent antibodies to determine cell phenotype. Cell sorting was conducted on a Cytek Aurora sorter. Sorted cells were preserved in TRIzol. mRNA isolation from TRIzol and cDNA synthesis were conducted using a standard protocol. Quantitative RT-PCR was conducted using *IKZF2* (Helios) specific primers 5’- ACGTGTGACAATGAGCTTTCA-3’ (forward) and 5’-ACTGAATTTGTGCTTGTCATGTG-3’ (reverse). SYBR green qPCR mastermix from Qiagen was used for qPCR. Relative units of mRNA were calculated against the expression of the housekeeping gene, *RPL13A*.

### Statistics

Student t tests or Mann–Whitney tests were conducted for single comparisons and ANOVA was conducted for multiple group analyses. When analyzing parametric single comparisons with unequal variances, t tests with Welch’s correction were used. When analyzing nonparametric multiple groups, Kruskal–Wallis tests were performed with false discovery rate adjustment.

## Results

### Short duration of HP infection in combination with autoimmunity sufficed to cause increase in gastric tumorigenesis, with loss of E-Cadherin and DMBT1 in gastric epithelial cells

CTLA4KD mice on the Balb/c x C57BL/6 F1 (CB6F1) background develop autoimmune gastritis and severe metaplastic lesions which spontaneously transition to invasive gastric adenocarcinoma after one year of age^13^. We took advantage of the full penetrance of tumorigenesis with clear landmarks of histopathological progression to test the combinatorial effect of autoimmunity and HP infection. Wildtype (WT) and CTLA4KD mice were infected with HP at 1-3 months of age and sacrificed at 9-10 months of age (WT HP and CTLA4KD HP, respectively). One-week post-infection HP was undetectable in the stool of most CTLA4KD HP mice, unlike in WT HP stool (Fig. 1A), probably due to autoimmune inflammation in the gastric mucosa limiting HP colonization. The results indicate that HP infection in the context of autoimmune gastritis in CTLA4KD HP mice persisted less than a week. At the endpoint, WT HP mice had increased HP CFU compared to CTLA4KD mice (Fig. 1B) and positive urease tests were observed in WT HP mice, but not CTLA4KD HP mice (Fig. S1A).

**Figure 1.**
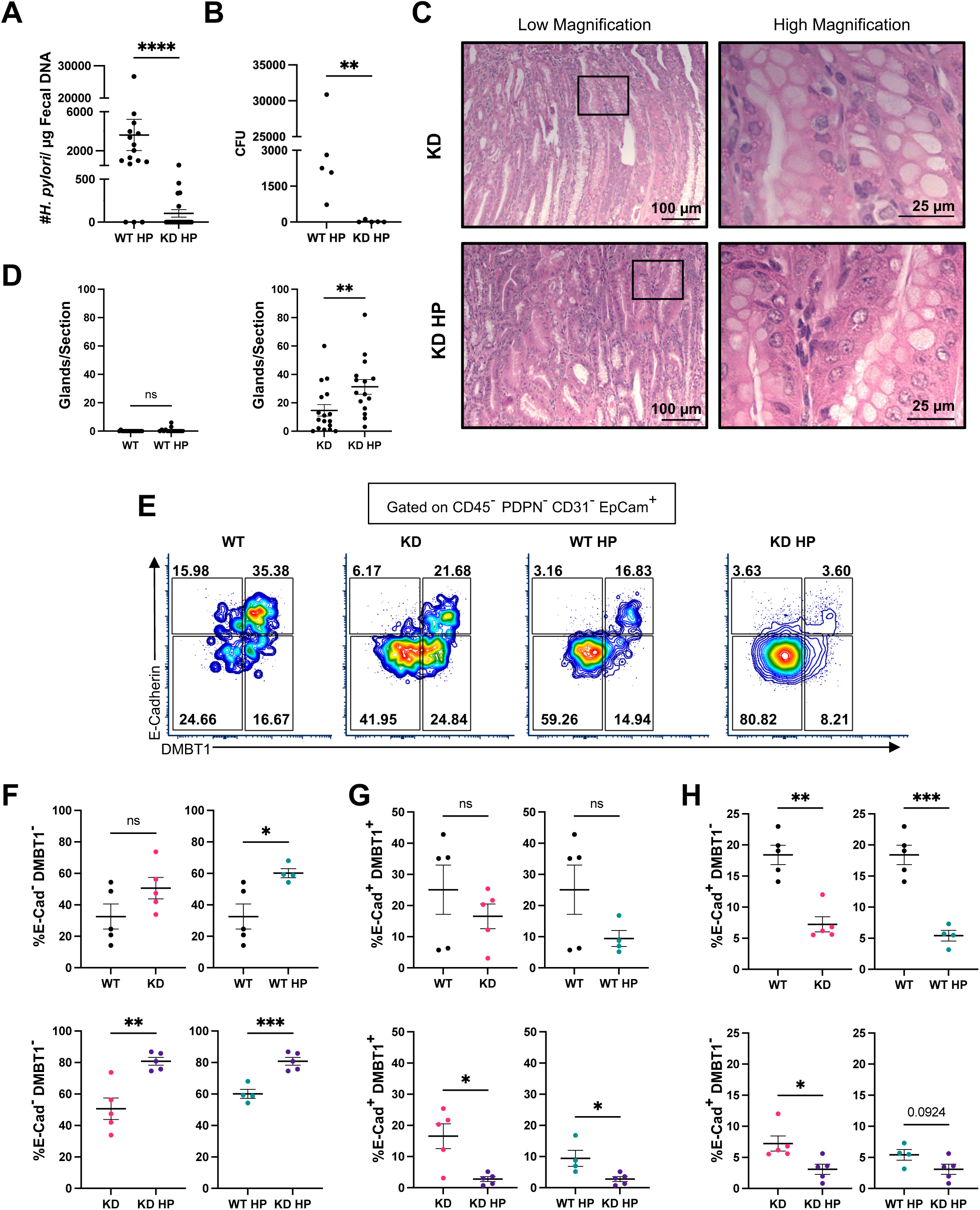
Effects of combinatorial autoimmunity and HP infection on the gastric mucosa and tumorigenic markers. WT and CTLA4KD mice on the CB6F1 genetic background were administered HP at 1-3 months of age and sacrificed 6-8 months post HP infection. Stomachs were analyzed by histopathology or enzymatically processed for single cell preparations for flow cytometry. (A) HP load within the stool of WT HP mice and CTLA4KD HP mice at 1 week post infection. (B) HP CFU counts from stomach homogenate cultures in WT HP and KD HP mice. (C) Representative low and high magnification images of gastric mucosa glands from the corpus presenting with globoid dysplasia. (D) Quantification of globoid dysplastic glands per stomach section in WT and WT HP mice (left) and in CTLA4KD and CTLA4KD HP mice (right). (E) Representative flow cytometry plots for E-Cadherin (E-cad) and DMBT1 expression in gastric epithelial cells. (F) Proportion of E-Cad^-^DMBT1^-^ cells in the stomach mucosa. (G) Proportion of E-Cad^+^DMBT1^+^ cells in the stomach mucosa. (H) Proportion of E-Cad^+^DMBT1^-^ cells in the stomach mucosa. Each data point represents one animal (Mean ± SEM; *p ≤ 0.05, **p ≤ 0.01, ***p ≤ 0.001. ****p ≤ 0.0001).

In this model, due to rampant inflammatory and metaplastic pathology caused by autoimmunity, we could not observe statistically significant changes in gross pathology impacted by the short duration of HP infection in the CTLA4KD group. However, we detected an increase in globloid dysplasia^19^ of the gastric mucosa in CTLA4KD HP mice when compared to CTLA4KD mice; no difference was found between WT HP and WT mice (Fig. 1C, 1D). Thus, despite the inability of HP to establish lasting infection in CTLA4KD HP mice, an additive effect between autoimmunity and HP infection was observed in the promotion of dysplastic changes within the gastric mucosa.

We characterized cellular profiles of the gastric mucosae with a 40-color spectral flow cytometry panel that allowed us to analyze major lineages in the nonhematopoietic and hematopoietic compartments. We examined epithelial cells characterized as CD45^-^ PDPN^-^CD31^-^ EpCam^+^ cells. HP infection reduced characteristic epithelial cell proteins, E-Cadherin, known for its association with GC development due to loss of function mutations^20^, and Deleted in Malignant Brain Tumors 1 (DMBT1), a putative tumor suppressor gene^21^ (Fig. 1E). A higher proportion of WT HP and CTLA4KD HP cells were E-Cadherin^-^DMBT1^-^, with an additive effect between autoimmunity and HP infection (Fig. 1F). In parallel, the E-Cadherin^+^DMBT1^+^ population was decreased in CTLA4KD HP mice compared to WT HP and CTLA4KD mice (Fig. 1G). HP infection alone reduced the population of E-Cadherin^+^DMBT1^-^ cells within the gastric epithelium, with a further reduction in the context of autoimmunity (Fig. 1H). No differences were observed in the proportion of E-Cadherin^-^DMBT1^+^ cells (Fig. S1B). These results indicate that autoimmunity in combination with HP infection, even for a short duration, advances tumorigenic differentiation of epithelial cells.

### Autoimmunity and HP infection collaboratively promote gastric mucosal inflammation in the form of tertiary lymphoid structures

Autoimmunity driven tumorigenesis in CTLA4KD mice is mediated by chronic type 2 inflammation^13^. Chronic inflammation is also a major mechanism in HP-driven GC development^14^. In non-lymphoid tissue, the formation of a tertiary lymphoid structure (TLS) sustains chronic inflammation^22^. We examined the combinatorial effect of autoimmunity and HP-infection on TLSs, using CTLA4KD mice on the C57BL/6 (B6) background, which does not permit full penetrance of autoimmune inflammation within the gastric mucosa. WT and CTLA4KD mice were infected with HP at ∼4 weeks of age and sacrificed at ∼3 months of age. In CTLA4KD HP and WT HP mice, HP was detected in stool one-week post infection (Fig. 2A). At the endpoint, urease tests revealed WT HP and CTLA4KD HP mice were colonized by HP (Fig. S2A) and CFU count showed no differences in colonization (Fig. 2B).

**Figure 2.**
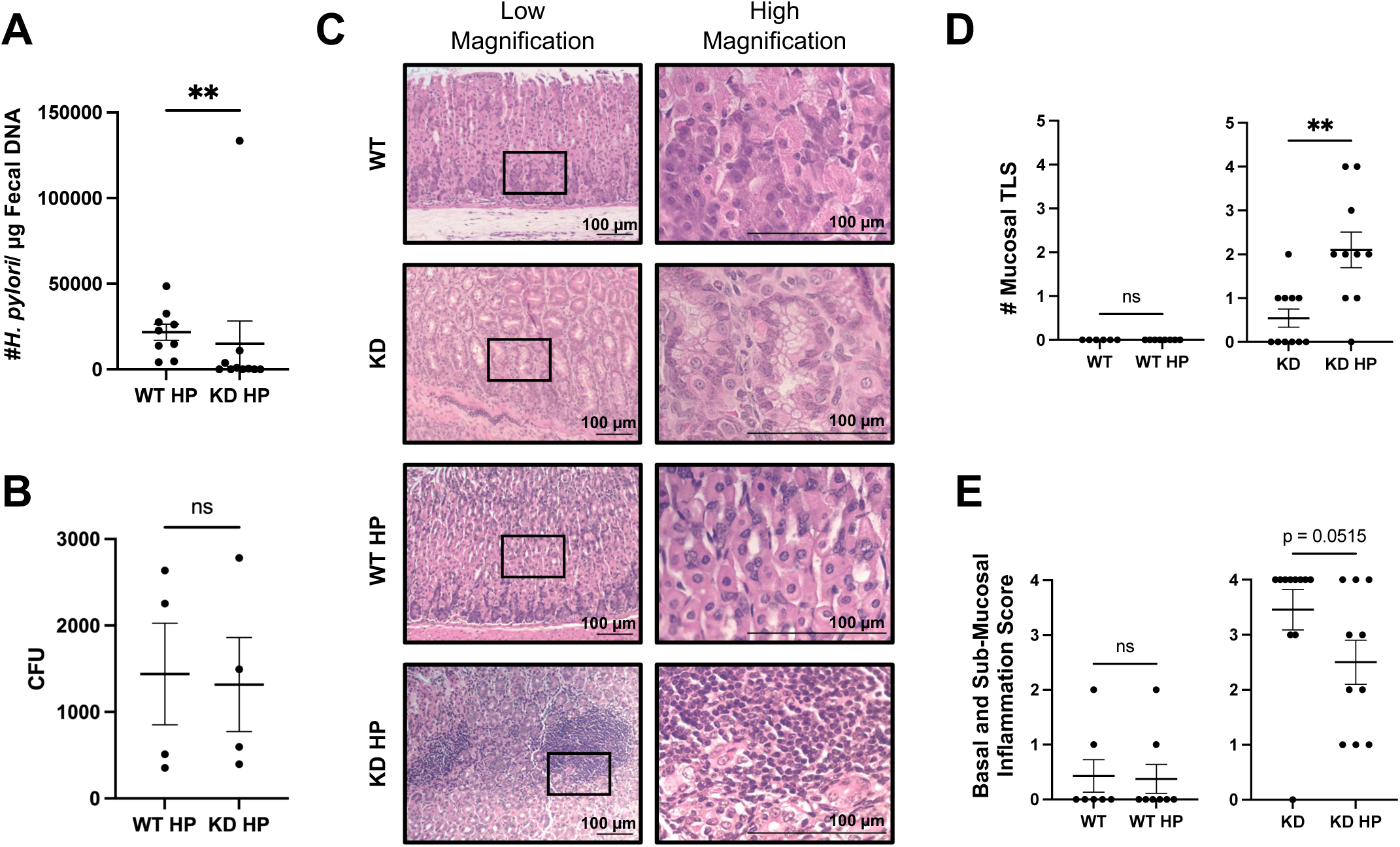
Effects of HP infection in combination with autoimmunity on chronic inflammation within the gastric mucosa. WT and CTLA4KD mice on the B6 genetic background were administered HP at ∼4 weeks of age and sacrificed 2-3 months post HP infection. (A) HP load within the stool of WT HP mice and CTLA4KD HP mice one week post infection. (B) HP CFU counts from stomach homogenate cultures in WT HP and KD HP mice.(C) Representative low and high magnification images of gastric mucosa glands and TLS in the corpus. Scale bar: 100 μm. (D) TLSs within the gastric mucosa. (E) Histopathology score of basal and sub-mucosal inflammation. Each data point represents one animal (Mean ± SEM; *p ≤ 0.05, **p ≤ 0.01)

Histopathological analysis of the stomach revealed that CTLA4KD HP mice had more TLSs compared to CTLA4KD mice (Fig. 2C-2D). Concurrently, CTLA4KD HP mice had reduced basal and sub-mucosal inflammation scores (Fig. 2E). While CTLA4KD mice can present with TLSs, inflammation is typically limited to the basal and sub-mucosal areas (Fig. 2C), suggesting that HP infection can alter the location of inflammatory infiltrates in the setting of autoimmunity to facilitate the proximal interaction between immune cells and mucosal epithelial lineages.

### The combination of HP infection and autoimmunity led to innate immune modulation

The coevolution of HP and humans likely reflects host immunosuppressive adaptations enabling HP persistence. Similarly, autoimmunity may trigger regulatory mechanisms that counter damage caused by inflammation. We first examined the combinatorial effect of HP and autoimmunity on innate immunity, the first line of immune defense against infection. We utilized dimensionality reducing viSNE analysis of 40-color spectral flow cytometry data from the spleen. Then we utilized the SPADE algorithm to further cluster cells and overlaid results on tSNE plots developed from viSNE analysis, allowing us to use relatively unbiased algorithms to cluster cell populations based on expression of defined markers. NK cells were found to be reduced in CTLA4KD HP mice (Fig. 3A). Manual gating of CD45^+^ CD19^-^ TCRβ^-^ NK1.1^+^ cells within the spleen demonstrated that HP infection reduced the proportion of splenic NK cells in WT HP mice, but not cell numbers (Fig. 3B-3D). In contrast, both NK cell proportions and numbers were reduced in CTLA4KD HP mice compared to CTLA4KD mice (Fig. 3B-3D). Autoimmunity and HP individually caused NK cell reductions, but CTLA4KD HP mice had levels of NK cells lower than the reductions caused by either alone (Fig, 3B-3D), indicating that autoimmunity and HP infection have an additive effect in reducing NK cells.

**Figure 3.**
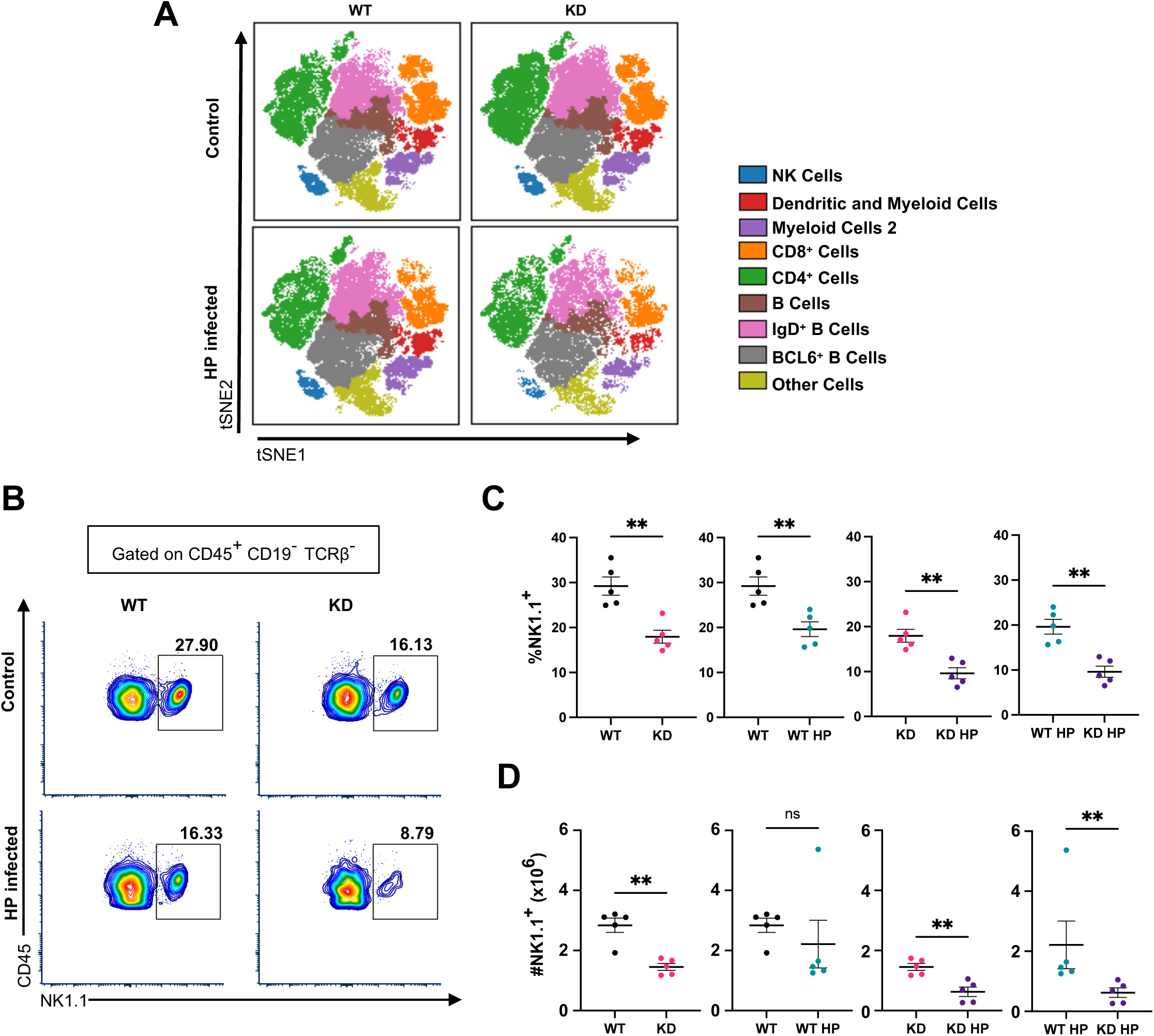
Systemic effect on innate immune response by autoimmunity and HP infection in spleen. The spleens of WT and CTLA4KD mice on the B6 background were analyzed 2-3 months post HP infection by flow cytometry. (A) Hematopoietic cell populations (gated on CD45^+^ cells, excluding dead cells and doublets) were examined by SPADE on viSNE analysis. Plots displaying the indicated cell populations. (B) Representative flow cytometry plots of NK1.1^+^ cells. (C) Proportion of NK1.1^+^ cells. (D) NK1.1^+^ cell numbers. Each data point represents one animal (Mean ± SEM, *p ≤ 0.05, **p ≤ 0.01)

### HP infection and autoimmunity upregulated the transcription factor Helios which promotes suppressive T cell activity

We initially hypothesized that HP infection promotes immunosuppression by CD4 T regulatory (CD4^+^ Foxp3^+^; Treg) cells, a major mechanism of peripheral tolerance^23^. We analyzed T cell profiles in GLN, MLN and SPL and did not detect change in the Treg population but found increased CD4 conventional (CD4^+^ Foxp3^-^; CD4 Tconv) cells expressing Helios. Helios, a transcriptional factor known for its role in stabilizing the Treg lineage, was originally believed to be exclusively expressed in thymic derived Tregs^24^. The expansion of Helios^+^ CD4 Tconv cells occurred not only locally (in the GLN) (Fig.4A, 4B, 4C), but also systemically in the MLN and SPL (Fig. S3A-S3F).

**Figure 4.**
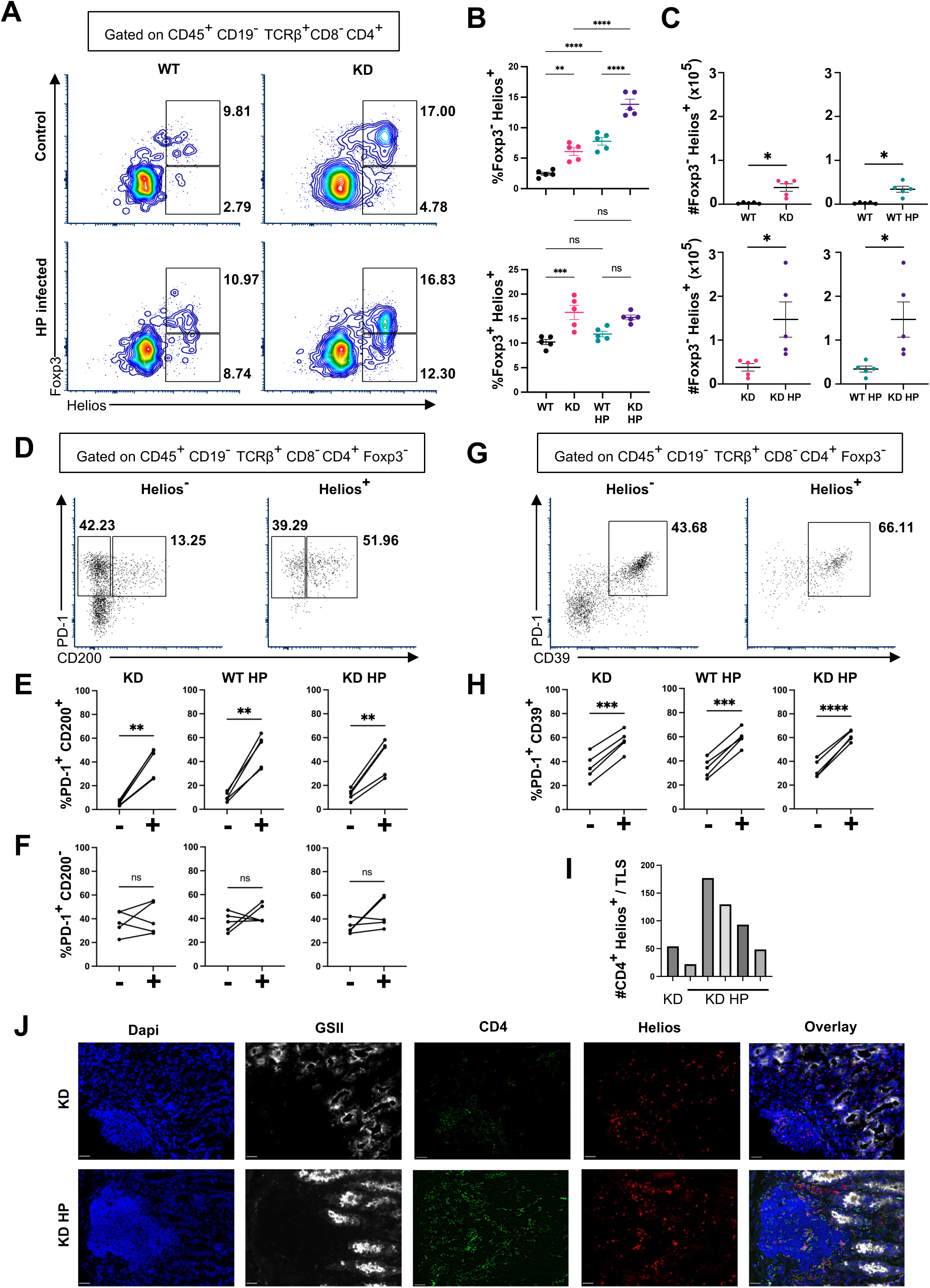
Expression of Helios in CD4 T cells in lymphoid organs and gastric TLS. The GLN from WT and CTLA4KD mice on the B6 background were analyzed 2-3 months post HP infection by flow cytometry. Stomach cryosections were analyzed by immunofluorescence. (A) Representative flow cytometry plots of Helios and Foxp3 expression in CD4^+^ T cells. (B) Percentage of cells expressing Helios in Foxp3^-^ (top) and Foxp3^+^ (bottom) cells. (C) Number of Foxp3^-^ Helios^+^ cells. (D) Representative flow cytometry plots of PD-1 and CD200 on Helios^-^ and Helios^+^ cells. (E) Percentage of Helios^-^ (-) and Helios^+^ (+) PD-1^+^ CD200^+^ cells. (F) Percentage of Helios^-^ (-) and Helios^+^ (+) PD-1^+^ CD200^-^ cells. (G) Representative flow cytometry plots of PD-1 and CD39 on Helios^-^ and Helios^+^ cells. (H) Percentage of Helios^-^ (-) and Helios^+^ (+) PD-1^+^ CD39^+^ cells. (I) Quantification of CD4^+^Helios^+^ cells per TLS per mouse from immunofluorescence staining. (J) Representative immunofluorescence images of a TLS from a CTLA4KD (top) and CTLA4KD HP (bottom) mouse in the stomach mucosa. Scale bar: 25 μm. Each data point represents one animal (Mean ± SEM, *p ≤ 0.05, **p ≤ 0.01, ***p ≤ 0.001. ****p ≤ 0.0001).

Helios^+^ CD4 Tconv cells were enriched for immune inhibitory markers PD-1 and CD200, suggesting their suppressive potentials. As shown in Fig. 4D, 4E, Helios^+^ CD4 Tconv cells co-express PD-1 and CD200 on their surface at a higher level than Helios^-^ CD4 Tconv cells. PD-1^+^CD200^-^ cells remained unchanged between Helios^-^ and Helios^+^ CD4 Tconv cells, suggesting upregulation of CD200 on PD-1^+^Helios^+^ CD4 Tconv cells (Fig. 4F). Expression of CD200, a critical immune checkpoint^25^, indicates Helios^+^ CD4 Tconv cells could dampen the response of other immune cells. Helios^+^ CD4 Tconv cells also displayed increased co-expression of PD-1 and CD39 (Fig. 4G, 4H). PD-1 expression on its own indicates T cell activation, but co-expression with CD39 suggests that CD4 cells may be exhausted and dysfunctional^26^.

Lastly, we measured CD4^+^Helios^+^ cells in gastric mucosal TLSs by immunofluorescence. TLSs were identified using dapi and CD4 staining, and an absence of GSII lectin binding, which stains gastric glands (Fig. 4J). Quantification of CD4^+^Helios^+^ cells in each TLS revealed that CTLA4KD HP mice had more CD4^+^Helios^+^ cells present in the gastric mucosa (Fig. 4I, 4J).

Results show that autoimmunity and HP interact to systemically elevate an immunosuppressive and tolerogenic CD4 Tconv profile by induction of Helios. Lastly, increased CD4^+^Helios^+^ cells in CTLA4KD HP gastric mucosa implies HP may utilize Helios to create a suppressive and tolerogenic inflammatory environment and establish chronic infection.

### HP infection led to long lasting Helios induction in CD4 cells

The CB6F1 model allowed us to follow the long lasting effects of HP infection in the context of autoimmunity. The GLN, MLN and SPL underwent analysis using the 40-color spectral flow cytometry panel. SPADE results from the GLN revealed a population of CD4^+^ Helios^+^ cells (Fig. 5A). Results were corroborated using manual gating. Within the GLN no difference in CD4 Tconv cells expressing Helios was detected in WT HP mice versus controls, but CTLA4KD HP mice displayed an increase in proportion and number (Fig. 5B, 5C, S4A). CTLA4KD HP GLNs had elevated numbers of Helios^+^ Tregs, but not in proportion (Fig. 5D, S4B). Within the MLN, CTLA4KD HP mice had an increased proportion of Helios^+^ CD4 Tconv cells (Fig. S4C-S4G). In the spleen, both WT HP and CTLA4KD HP had increases in Helios^+^ CD4 Tconv cell proportions and numbers when compared to non-infected controls (Fig. S4H-S4L). Helios^+^ CD4 Tconv cells were also found to co-express PD-1 with CD200 and PD-1 with CD39 at higher levels than Helios^-^ CD4 Tconv cells in GLN, MLN and SPL (Fig. 5E-5I, S5A-S5H). These results suggest that HP infection in combination with autoimmunity leads to long-lasting systemic alteration of Helios^+^ CD4 Tconv cells expressing inhibitory and exhaustion markers.

**Figure 5.**
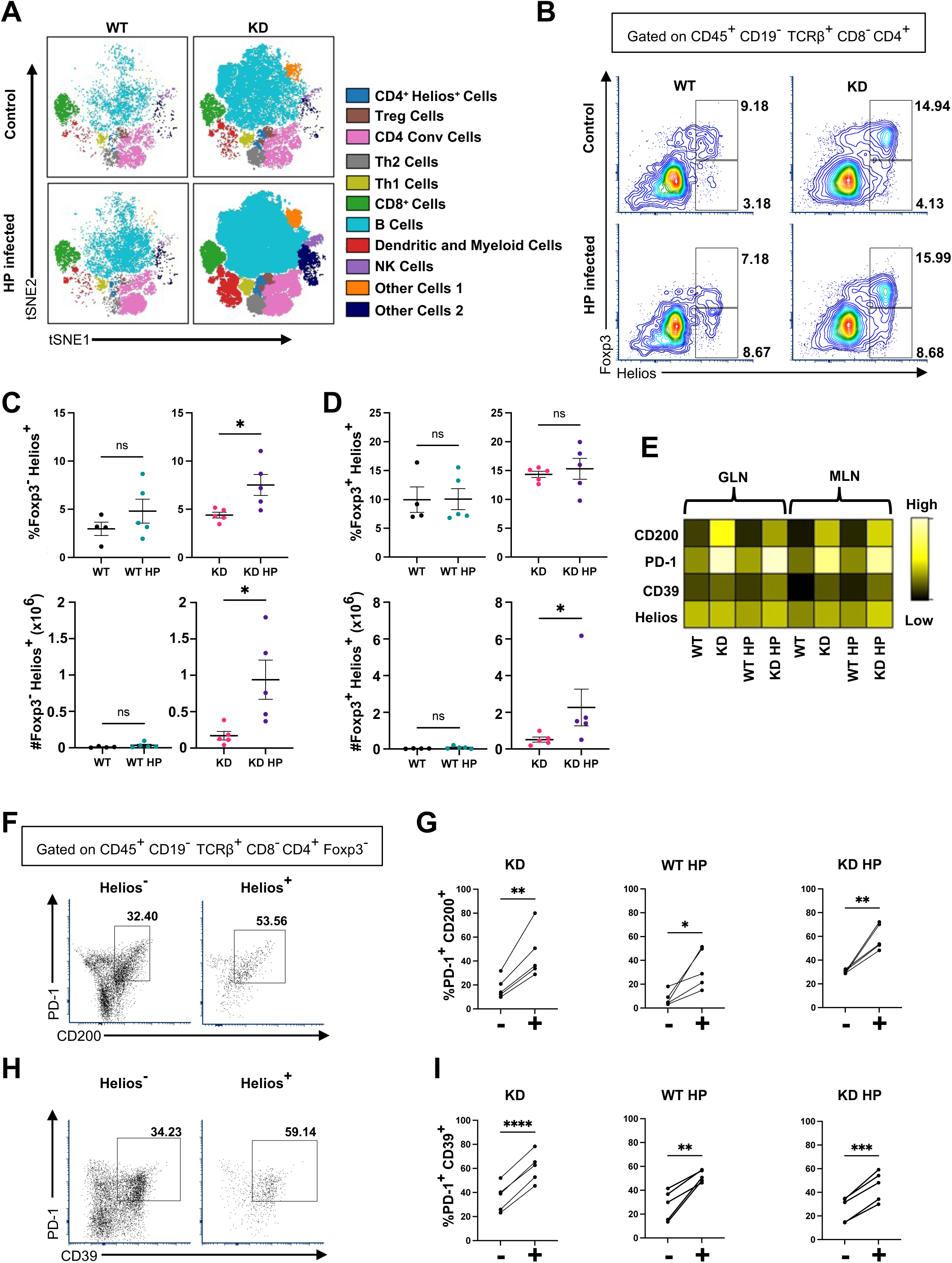
Lasting effect of HP infection on Helios induction and upregulation of immune suppression in the setting of autoimmunity. The lymph nodes from WT and CTLA4KD mice on the CB6F1 background were analyzed 6-8 months post HP infection by flow cytometry. (A) Spade on viSNE analysis of GLN displaying cell populations indicated on the right. (B) Representative flow cytometry plots showing Foxp3 and Helios expression in CD4 cells from the GLN. (C) Foxp3^-^Helios^+^ cell percentages (top) and cell counts (bottom) from the GLN. (D) Foxp3^+^Helios^+^ cell percentages (top) and cell counts (bottom) from the GLN. (E) Spade analysis of Helios, CD39, PD-1 and CD200 in CD4 cells from the GLN and MLN. (F) Representative flow cytometry plots showing expression of PD-1 and CD200 in Helios^-^ and Helios^+^ cells from the GLN. (G) Percentage of Helios^-^ (-) and Helios^+^ (+) PD-1^+^ CD200^+^ cells from the GLN. (H) Representative flow cytometry plots showing expression of PD-1 and CD39 in Helios^-^ cells and Helios^+^ cells from the GLN. (I) Percentage of Helios^-^ (-) and Helios^+^ (+) PD-1^+^ CD39^+^cells from the GLN. (F-I) Gated on CD45^+^CD19^-^TCRβ^+^CD8^-^CD4^+^Foxp3^-^ cells. Each data point represents one animal (Mean ± SEM, *p ≤ 0.05, **p ≤ 0.01, ***p ≤ 0.001. ****p ≤ 0.0001.)

### HP infection led to systemic modulation of CD8 T cell phenotype

We next analyzed the CD8 T cell compartment to examine Helios expression. The proportion of CD8 Helios^+^ cells was elevated in WT HP SPL when compared to WT SPL. No difference was detected between CTLA4KD and CTLA4KD HP groups (Fig 6A, 6B). Despite the increased proportion of CD8 Helios^+^ cells in WT HP SPL, no difference in numbers was found when compared to WT SPL (Fig. 6C). Within the GLN there was an increase in proportion of CD8 Helios^+^ cells in WT HP compared to WT mice, but no difference in numbers (Fig. S6A-S6B). No changes were observed in the MLN (Fig. S6C-S6D). These results suggest that HP can systemically induce Helios expression in CD8 cells, along with a local effect in the GLN.

**Figure 6.**
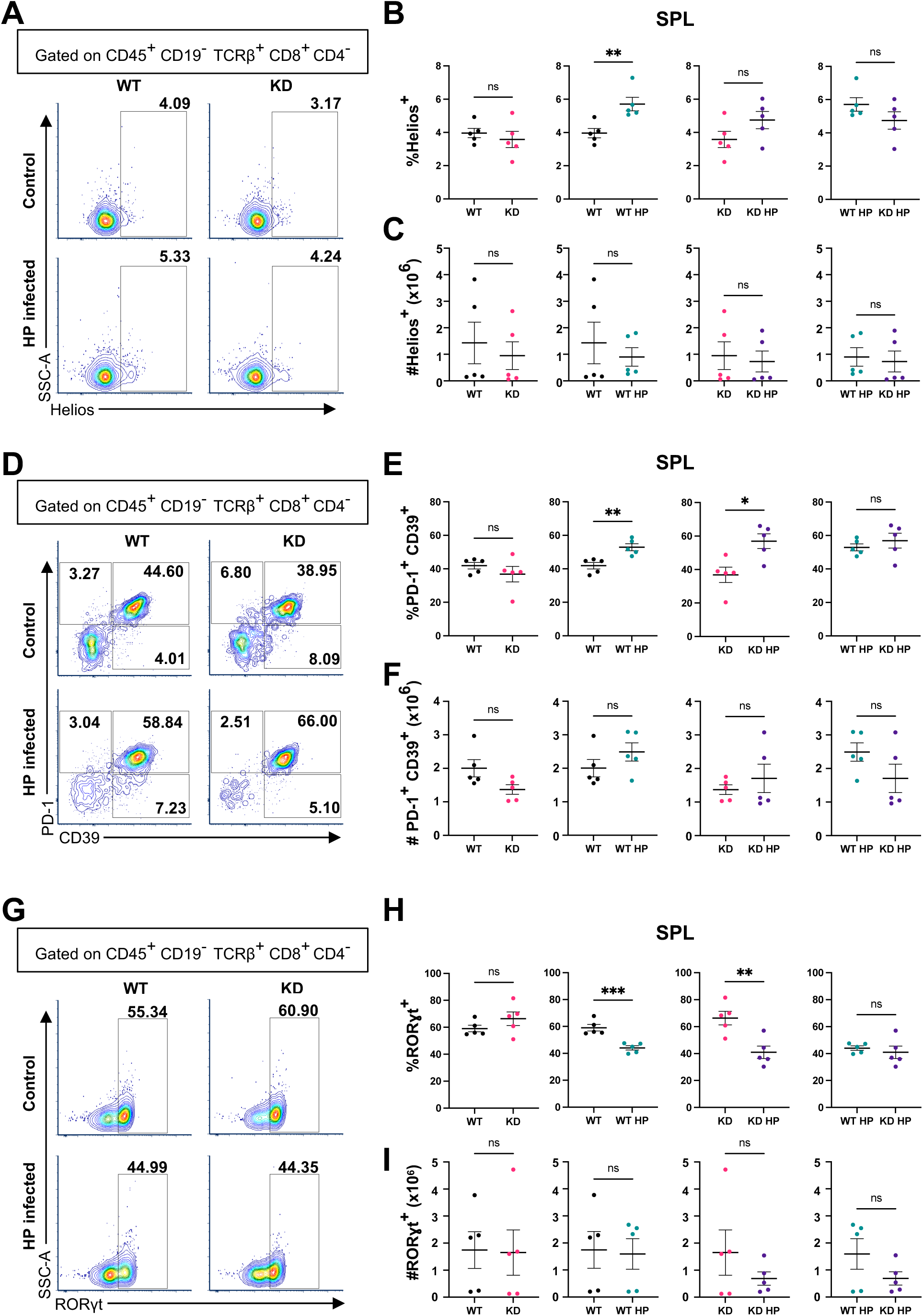
Systemic effect on CD8 T cells from HP infection and autoimmunity. The spleens from WT and CTLA4KD mice on the B6 background were analyzed 2-3 months post HP infection by flow cytometry. (A) Representative flow cytometry plots of Helios expression in CD8 T cells. (B) Percentage of CD8 T cells expressing Helios. (C) Number of CD8^+^Helios^+^ cells. (D) Representative flow cytometry plots of PD-1 and CD39 expression on CD8 T cells. (E) Proportion of CD8 T cells co-expressing PD-1 and CD39. (F) Number of PD-1^+^CD39^+^ CD8 T cells. (G) Representative flow cytometry plots of RORγt expression in CD8 T cells. (H) Percentage of CD8 T cells expressing RORγt. (I) Number of RORγt^+^ cells. Each data point represents one animal (Mean ± SEM, n = 5, *p ≤ 0.05, **p ≤ 0.01, ***p ≤ 0.001.)

Work from others found that Helios in CD8 cells coincides with exhaustion and limited effector function^27^. Phenotypic analysis of CD8 cells, regardless of Helios expression, revealed an increased proportion of PD-1^+^CD39^+^ cells in response to HP (Fig. 6D, 6E). No difference in cell numbers was observed (Fig. 6F). These results were unique to the spleen, with no difference in the GLN or MLN (Fig S6E-S6H), suggesting a systemic effect

Expression of the transcription factor RORγt (Retinoic acid-related orphan receptor gamma t) in CD8 T cells is associated with a proinflammatory response^28^. We analyzed RORγt in CD8 T cells. In response to HP infection, the proportion of splenic CD8 T cells expressing RORγt decreased (Fig. 6G, 6H), while numbers remained unaltered (Fig. 6I), likely due to variation within groups. No difference in CD8^+^RORγt^+^ cells was present in the GLN or MLN (Fig. S6I-S6L). Downregulation of RORγt is likely to dampen the immune response due to the association of CD8^+^ RORγt^+^ cells with a proinflammatory response. Overall, these results indicate that HP has a strong systemic effect on CD8 T cell effector function.

CB6F1 mice were used to analyze the effect of long-term HP infection on CD8 cells. Results were similar to those observed in B6 short-term HP mice. The GLN did not demonstrate changes in Helios expression in WT HP or CTLA4KD HP mice (Fig. S7A, S7B). Within the MLN, HP infection caused increased CD8^+^Helios^+^ cells in WT HP and CTLA4KD HP mice when compared to WT and CTLA4KD mice, respectively (Fig. S7C, S7D). Lastly, splenic CD8^+^Helios^+^ cells were elevated in CTLA4KD HP mice compared to CTLA4KD mice (Fig. S7E, S7F). We also analyzed expression of PD-1 and CD39 on CD8 T cells to assess exhaustion and found alterations to PD-1 and CD39 expression in the GLN are retained long term after HP clearance (Fig. S6G-S6I). Results indicate that similar to CD4 cells, CD8 T cells from CTLA4KD HP mice undergo systemic expansion of CD8^+^Helios^+^ cells and exhaustion markers following HP infection that is preserved long-term after clearance.

### Human PBMCs upregulate Helios in CD4 cells *in vitro* in response to HP co-culture

Conceivably, HP infection in humans may lead to direct or proximal interaction between HP and immune cells in gastric mucosae, a process difficult to model. We used an *in vitro* co-culture model for initial assessment into whether Helios expression in T cells can be affected directly by HP. Stimulated human PBMCs co-cultured with HP or heat killed HP at a MOI of 2 for 16 hours were analyzed for Helios protein expression. Co-culture of HP led to reduction of both CD4 and CD8 T cells (Fig 7A). Due to the drastic decrease observed in CD4 T cells there are reduced numbers of both CD4^+^CD25^+^CD127^+^ and CD4^+^CD25^+^CD127^-^ that are Helios^+^ in HP co-cultured samples (Fig. 7B,7C). Higher proportions of CD4^+^CD25^+^CD127^+^ cells (conventional CD4 cells) expressed Helios when compared to heat killed HP controls. CD4^+^CD25^+^CD127^-^(enriched for Treg) cells from the same donors did not display increased Helios expression in HP co-cultured samples (Fig. 7B, 7D).

**Figure 7.**
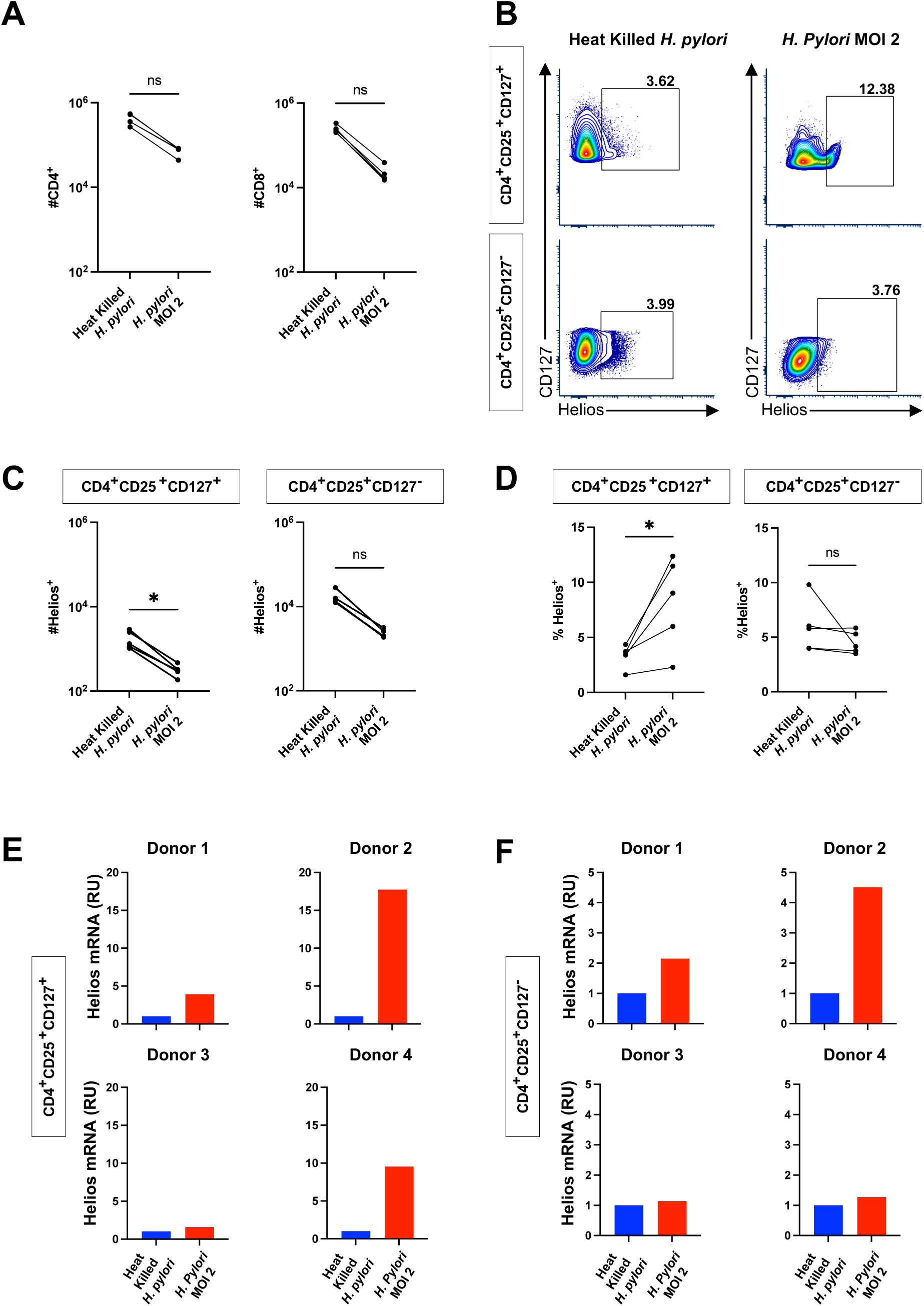
Helios expression in Human CD4 T cells in response to HP infection *in vitro*. Human donor (n=4) PBMCs were stimulated and co-cultured with either heat killed *H. pylori* or *H. pylori* at an MOI of 2 for 16 hours. Cells were then analyzed by flow cytometry or underwent cell sorting followed by qRT-PCR. (A) Number of CD4 T cells (left) and CD8 T cells (right). Cell numbers are normalized to per million total cells. (B) Representative flow cytometry plots displaying expression of CD127 and Helios on stimulated CD4 T Cells. (C) Number of CD4^+^CD25^+^CD127^+^ and CD4^+^CD25^+^CD127^-^ cells expressing Helios. Cell numbers are normalized to per million total cells. (D) Proportion of CD4^+^CD25^+^CD127^+^ and CD4^+^CD25^+^CD127^-^ cells expressing Helios (*p ≤ 0.05). (E-F) mRNA expression of Helios was measured in (E) CD4^+^ CD127^+^CD25^+^ cells and (F) CD4^+^ CD127^-^CD25^+^ cells.

To examine whether increased proportions of Helios^+^ cells merely reflect preferential survival or are contributed by induction of Helios expression, we sorted the cell subsets and measured Helios mRNA. Activated human PBMCs were co-cultured with HP at an MOI of 2 or heat killed HP for 16 hours and then sorted for CD4^+^CD25^+^CD127^+^ and CD4^+^CD25^+^CD127^-^cells. Helios mRNA was measured by qRT-PCR in reference to house-keeping gene expression. All four donors had increased Helios mRNA in CD4^+^CD25^+^CD127^+^ cells when compared to heat killed HP controls (Fig. 7E). CD4^+^CD25^+^CD127^-^ cells also displayed an increase in Helios mRNA (Fig. 7F).

Co-culture of PBMCs with HP is an artificial model of HP and immune cell interaction. Nevertheless, PBMC coculture with HP provided insights into the potential effect from the direct or proximal interaction of HP and T cells. The *in vitro* upregulation of Helios mRNA and protein in CD4 Tconv in the human cells is consistent with the findings from the *in vivo* murine models.

## Discussion

In this study we tested a hypothesis that HP infection and autoimmunity exert a combinatorial effect on stomach tumorigenesis. Our findings indicate that HP infection and autoimmunity collaboratively promote tumorigenesis, affecting both the gastric mucosa and immune system. Combined HP infection and autoimmunity increased dysplastic changes within the gastric mucosa and promoted pro-tumorigenic differentiation of gastric epithelial cells. The immune landscape was also modified with reduced NK cells, induction of Helios in CD4 Tconv cells, and systemic modulation of CD8 cells.

As autoimmune diseases rise, it becomes likely that HP and gastric autoimmunity will converge in the same individual, as reported in a patient with CTLA4 haploinsufficiency^12^. Another case of an individual diagnosed with HP and autoimmune gastritis showed that atrophy rapidly advanced following eradication of HP^29^. Our study provides the first experimental evidence (to our knowledge) of the molecular and cellular interplay of HP infection and autoimmunity, offering mechanistic insight into how their convergence may accelerate GC development. Although the CTLA4KD model is built to mimic the genetic of GC in patients with CTLA4 haploinsufficiency, the inflammatory pathway and epigenetic modifications underlining GC development is broadly applicable to human GC development in general^13^.

Studies have shown that prolonged HP infection can initiate GC development, eventually leading to a decline or elimination of HP peristence^30^. Strikingly, even when pre-existing autoimmunity limited long-term persistence of HP infection, their combined effects on the gastric mucosa persisted for 6-8 months post infection, including increased globoid dysplasia and loss of tumor suppressors, E-Cadherin and DMBT1. Globoid (tubule neck) dysplasia is a precursor to diffuse type GCs, including signet ring cell carcinoma, which are frequently associated with *CDH1* mutations, the gene encoding E-Cadherin^31^. In addition, loss of E-Cadherin also marks epithelial-mesenchymal transition preceding metastasis^20^. Moreover, HP infection can induce E-Cadherin loss through CagA mediated disruption of the E-Cadherin/β-Catenin complex^32^. E-Cadherin downregulation has also been detected in response to inflammation^33, 34^.DMBT1 is a potential tumor suppressor found to play a role in gastric epithelial cell differentiation, with reduced expression in GC^35^. The downregulation of E-Cadherin and DMBT1 could be contributed by CagA and other virulence factors of HP as well as inflammatory effectors elicited from autoimmunity and/or HP infection. Although the exact mechanisms remains unknown for the additive effect between HP and autoimmunity in E-Cadherin and DMBT1 reduction, it is associated with increased globoid dysplasia of the gastric mucosa, and thus could foreseeably contribute to the development of early onset GC in younger populations.

The combination of HP and autoimmunity promoted TLSs in the gastric mucosa. HP can alter the location of inflammatory infiltrates, deeper into the mucosae, as opposed to the characteristic inflammation in CTLA4KD mice localized within the basal and sub-mucosal areas^13^. Lymphoid aggregates are a recognized symptom of HP infection and are present in ∼75% of gastric biopsies from HP infected individuals^36^. TLSs are a double-edged sword: they can promote anti-tumor immunity in cancer but may promote pathogenesis in autoimmune diseases^37^. In the complex environment of HP infection and autoimmunity, it is unknown whether TLSs play a pathogenic or protective role; further studies on their role in inflammatory tumorigenesis and/or antitumor immunity in the local environment are warranted.

Systemically HP and autoimmunity in combination present a striking effect on modulating innate and adaptive immunity. The effect on innate immunity is most pronounced in NK cell reduction. A decrease in NK cells has been detected in some autoimmune diseases^38^, such as systemic lupus erythematosus^39^ and type 1 diabetes^40^. In addition, research on HP found that infected individuals have a lower proportion of NK cells within the gastric mucosa^41^. The negative additive effect on NK cell numbers from autoimmunity and HP infection implies distinctive mechanisms mediated by the two causes.

Within the adaptive branch, our previous study demonstrated that autoimmune tumorigenesis in CTLA4KD mice is driven by Th2 cells^13^. Inflammation initiated in response to HP is also a major driver in the cascade to cancer. The immune response against HP is dominated by IFN-ɣ and Th1 cells^42^, although HP modulates the immune system towards a tolerogenic response driven by IL-10 and Tregs to establish chronic infection^26, 43^. Another study found the early response to HP is Th1 dominated followed by a Th2 response later^44^. Our results revealed expansion of Helios^+^ CD4 Tconv cells expressing inhibitory and exhaustion markers PD-1, CD39 and CD200 which may be a mechanism used by HP to induce tolerance in a pre-existing inflammatory environment. In addition, Helios^+^ CD4 Tconv cells remain elevated months post HP clearance in CTLA4KD mice. Studies have found that Helios can be induced in CD4 Tconv cells upon detection of autoantigen, leading to anergy^45^. Further functional studies on Helios^+^ CD4 Tconv cells and their response to HP and autoimmune stimulus are warranted to determine their role in tumorigenesis and early onset cancer, especially in clinical settings. With an *in vitro* culture model to capture the potential effect of proximal or direct interaction between HP and human immune cells, our findings revealed that CD4 Tconv cells from healthy human PBMCs upregulate Helios at the mRNA and protein levels in response to HP. Future studies of these cells and their long-term effects on GC development in humans are needed.

Unlike CD4 T cells, CD8 T cells, are not well-studied and not clearly understood in GC. An influx of CD8 T cells into the gastric mucosa occurs after HP infection and are later replaced by CD4 cells during chronic infection^46^. Our examination in the combinatorial setting found CD8 T cells were systemically altered, with pronounced alterations observed in the spleen where HP infection clearly increased Helios expression in CD8 T cells. Studies have shown that Helios expression in CD8 T cells limits effector function and may be a marker of regulatory CD8 T cells^27, 47^. Helios expression along with reduced RORγt and increased exhaustion markers strongly suggest downregulation of CD8 T cell function, but future studies into the role of CD8 Helios^+^ cells and CD8 T cell dysfunction in HP infection, autoimmunity and early onset GC is needed to understand their importance.

Overall, findings from this study provide initial insight into how HP and autoimmunity in combination may impinge lasting impacts through a multi-pronged effect on epithelial differentiation as well as systemic and local immune modulation. However, this study is limited in its scope and depth. It remains unknown if any HP virulence factors contribute to the lasting effect of HP in the setting of autoimmunity on innate and adaptive immune modulation. Furthermore, our current model is limited to a setting of pre-existing autoimmunity, as the sequence of HP colonization and autoimmunity onset may vary in humans and affect their interaction. Perhaps, mostly conspicuously, can lasting immune suppression be reversed? With a persistent effect, even after short HP exposure, additional remedies may be needed besides HP eradication, including interventions to reverse immune suppression. Taken together, this study provides new mechanistic insights and highlights the need for more comprehensive studies on the interaction of HP and autoimmunity, to understand the exact impact of HP and autoimmunity combination on early-onset GC, for the long-term goal of developing interventions against tumorigenic and immunosuppressive alterations to curtail and reverse the cascade of GC development.

## Abbreviations

GC: gastric cancer;
HP: *Helicobacter pylori*;
CTLA4KD: CTLA4 knockdown;
WT: wildtype;
SPEM: spasmolytic polypeptide expressing metaplasia;
GLN: gastric lymph nodes;
MLN: mesenteric lymph nodes;
SPL: spleen;
STO: stomach;
TLS: tertiary lymphoid structure;
CB6F1: Balb/c x C57BL/6 F1

## Authors’ Contributions

Emily N. Vazquez (conceptualization, formal analysis, investigation, methodology, writing- original draft, writing- review and editing), Clarisel Lozano (Investigation, writing- review and editing), Amanda M. Honan (methodology, writing- review and editing), Elena Zaika (investigation, resources), JeanMarie Houghton (conceptualization, writing- review and editing), Wael El-Rifai (conceptualization), Alexander Zaika (conceptualization, methodology, resources, writing- review and editing), Zhibin Chen (conceptualization, funding acquisition, methodology, resources, supervision, writing-review and editing)

## Data transparency statement

The data, methods used in the analysis, and materials used to conduct the research will be made available to any researcher for purposes of reproducing the results or replicating the procedure.

---

**Supplemental Table 1.** Antibodies utilized in 40-color spectral flow cytometry of murine gastric epithelial and lymphoid cells.

| **Antibody** | **Antibody Clone** | **Source** | **Identifier** |
| --- | --- | --- | --- |
| CD39 BUV395 | Y23-1185 | BD Biosciences | 567264 |
| CD11c BUV563 | HL3 | BD Biosciences | 749091 |
| PD-L1 BUV615 | MIH5 | BD Biosciences | 752339 |
| KLRG1 BUV661 | 2F1 | BD Biosciences | 741586 |
| CD19 BUV737 | 1D3 | BD Biosciences | 612781 |
| E-Cadherin BUV805 | DECMA-1 | BD Biosciences | 752480 |
| SOX2 BV421 | 14A6A34 | Biolegend | 656114 |
| CD4 Super Bright 436 | GK1.5 | Invitrogen | 62-0041-82 |
| Foxp3 eFluor450 | FJK-16s | Invitrogen | 48-5773-82 |
| Helios BV480 | 22F6 | Invitrogen | 414-9883-42 |
| Ki-67 BV510 | B56 | BD Biosciences | 563462 |
| TCRβ BV570 | H57-597 | Biolegend | 109231 |
| Ly-6A/3 (Sca-1) BV605 | D7 | Biolegend | 108134 |
| EpCam BV650 | G8.8 | Biolegend | 118241 |
| CD66a BV711 | CC1 | BD Biosciences | 750880 |
| CD11b BV750 | M1/70 | Biolegend | 101267 |
| CD200 BV786 | OX-90 | BD Biosciences | 744653 |
| MUC2 | HL1724 | Thermo Fisher | MA5-47157 |
| MHCII Alexa Fluor 532 | M5/114.15.2 | Invitrogen | 58-5321-82 |
| CD45 NovaFluor Blue 610-70s | 30-F11 | Invitrogen | M005T02B06 |
| CD31 NovaFluor Blue 660-120s | 390 | Invitrogen | M019T02B08 |
| IgD PerCP | 11-26c.2a | Biolegend | 405736 |
| RORγt PerCP eFluor710 | AFKJS-9 | Thermo Fisher | 46-6988-82 |
| Embigin PE | G7.43.1 | Invitrogen | 12-5839-82 |
| CD8a Spark YG593 | QA17A07 | Biolegend | 155024 |
| CD133 PE-Dazzle594 | 315-2C11 | Biolegend | 141212 |
| CD103 PE-FIRE640 | QA17A24 | Biolegend | 156918 |
| T-Bet PE-Cy5 | eBio4B10 (4B10) | Invitrogen | 15-5825-82 |
| CD38 PE-FIRE700 | 90 | Biolegend | 102748 |
| IL-4Rα PE-Cy7 | I015F8 | Biolegend | 144806 |
| PD-1 PE-FIRE810 | 29F.1A12 | Biolegend | 135253 |
| GLUT1 | SA0377 | Invitrogen | MA5-31960 |
| GATA3 eFluor660 | TWAJ | Thermo Fisher | 50-9966-42 |
| CD44 NovaFluor Red 700 | IM7 | Invitrogen | M010T02R03 |
| BCL6 R718 | K112-91 | BD Biosciences | 566979 |
| PDPN APC-FIRE750 | 8.1.1 | Biolegend | 127426 |
| DMBT1 Alexa Fluor 790 | G-4 | Santa-Cruz Biotechnology | Sc-514566 |
| NK1.1 APC-FIRE810 | S17016D | Biolegend | 156519 |

**Supplemental Figure 1.**
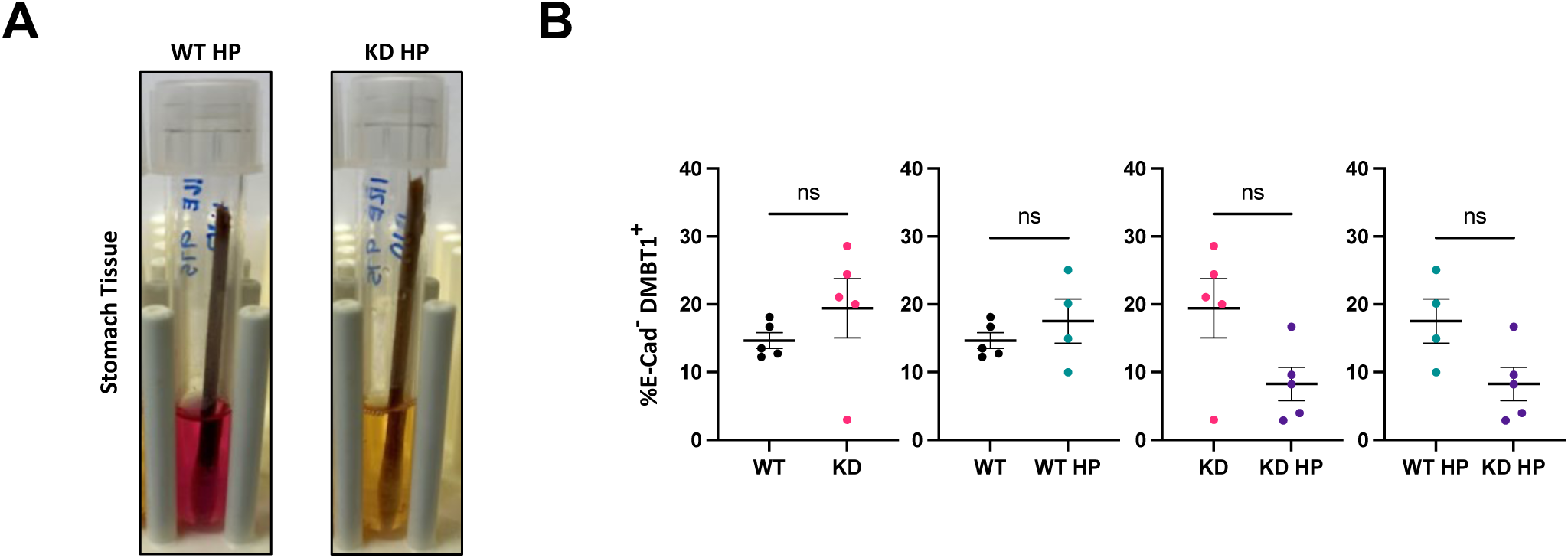
HP colonization and effect on proportion of E-Cadherin^-^DMBT1^+^ cells from the gastric mucosa. WT and CTLA4KD mice on the CB6F1 genetic background were administered HP by oral gavage at 1-3 months of age. Animals were sacrificed 6-8 months post HP infection. At the end point the stomach underwent enzymatic digestion and was analyzed by flow cytometry. (A) Representative images of urease test results from stomach homogenate cultures of HP infected animals at the end point (WT HP n=5, CTLA4KD HP n=5). (B) Proportion of cells that are E-Cad^-^DMBT1^+^ from the gastric mucosa. Each data point represents one animal (Mean± SEM, ns= not significant)

**Supplemental Figure 2.**
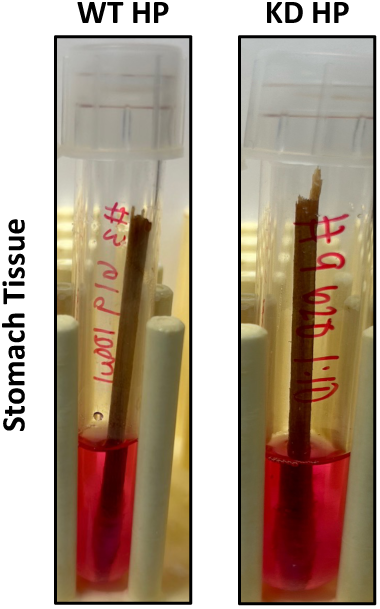
HP colonization of B6 gastric mucosa at the end point. Representative images of urease test results following culture of stomach homogenates at the end point from WT HP mice (n=4) and CTLA4KD HP mice (n=6).

**Supplemental Figure 3.**
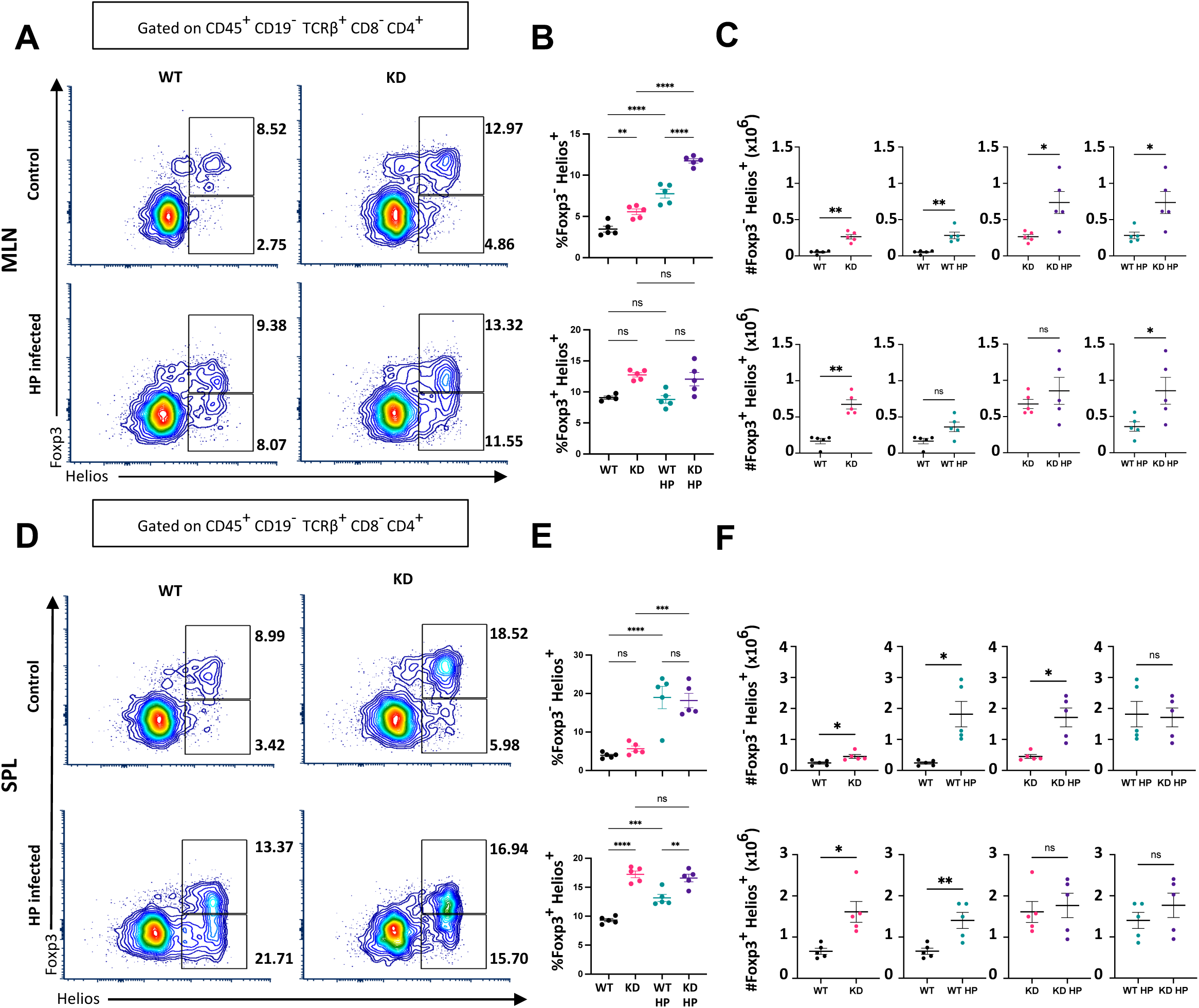
Induction of Helios in CD4 T cells in B6 MLN and SPL. WT and CTLA4KD mice on the B6 genetic background were administered HP by oral gavage at ∼4 weeks of age. Animals were sacrificed 2-3 months post HP infection. Single cell preparations were made from lymphoid organs and subsequently analyzed by flow cytometry. (A) Representative flow cytometry plots of Foxp3 and Helios expression in CD4^+^ T cells from the MLN. (B) Percentage of cells expressing Helios in Foxp3^-^ cells (top) and Foxp3^+^ cells (bottom) from the MLN. (C) Number of Foxp3^-^ Helios^+^ cells (top) and Foxp3^+^ Helios^+^ cells from the MLN. (D) Representative flow cytometry plots of Foxp3 and Helios expression in CD4^+^ T cells from the spleen (SPL). (E) Percentage of cells expressing Helios in Foxp3^-^ cells (top) and Foxp3^+^ cells (bottom) from the SPL. (F) Cell counts of Foxp3^-^Helios^+^ cells (top) and Foxp3^+^ Helios^+^ cells (bottom) from the SPL. Each data point represents one animal (Mean± SEM, *p ≤ 0.05, **p ≤ 0.01, ***p ≤ 0.001. ****p ≤ 0.0001)

**Supplemental Figure 4.**
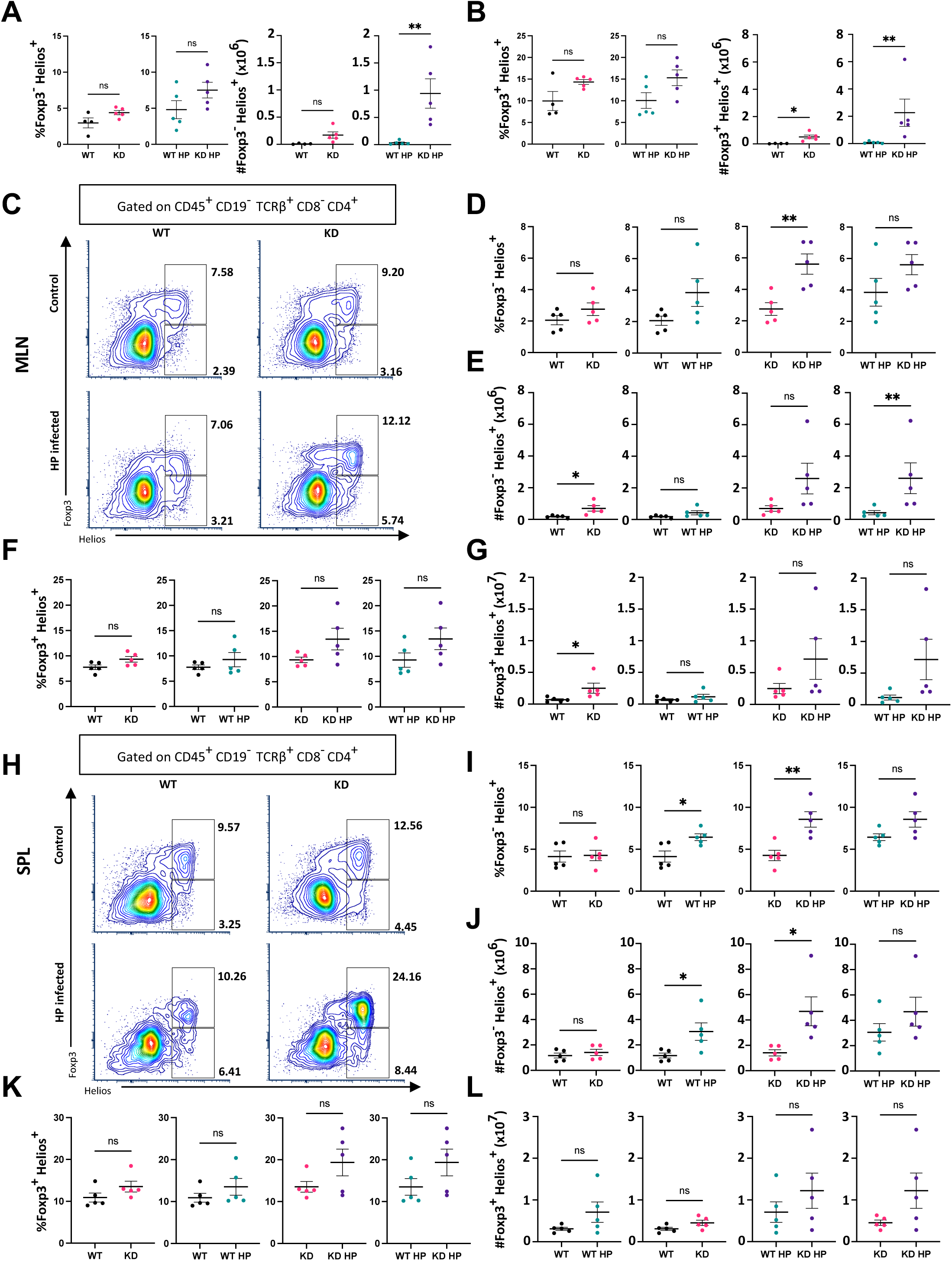
Systemic induction of Helios long-term. WT and CTLA4KD mice on the CB6F1 genetic background were administered HP by oral gavage at 1-3 months of age. Animals were sacrificed 6-8 months post HP infection. Single cell preparations were made from lymphoid organs and subsequently analyzed by flow cytometry. (A) Proportion (left) and cell number (right) of Foxp3^-^ Helios^+^ CD4 cells from the gastric lymph nodes (GLN). (B) Proportion (left) and cell number (right) of Foxp3^+^ Helios^+^ CD4 cells from the GLN. (C) Representative flow cytometry plots showing Foxp3 and Helios from the mesenteric lymph nodes (MLN). (D) Quantification of Foxp3^-^Helios^+^ cell percentages from the MLN. (E) Quantification of Foxp3^-^Helios^+^ cell counts from the MLN. (F) Quantification of Foxp3^+^Helios^+^ cell percentages from the MLN. (G) Quantification of Foxp3^+^Helios^+^ cell counts from the MLN. (H) Representative flow cytometry plots showing Foxp3 and Helios from the SPL. (I) Quantification of Foxp3^-^Helios^+^ cell percentages from the SPL. (J) Quantification of Foxp3^-^Helios^+^ cell counts from the SPL. (K) Quantification of Foxp3^+^Helios^+^ cell percentages from the SPL. (L) Quantification of Foxp3^+^Helios^+^ cell counts from the SPL. Each data point represents one animal (Mean ± SEM, *p ≤ 0.05, **p ≤ 0.01.)

**Supplemental Figure 5.**
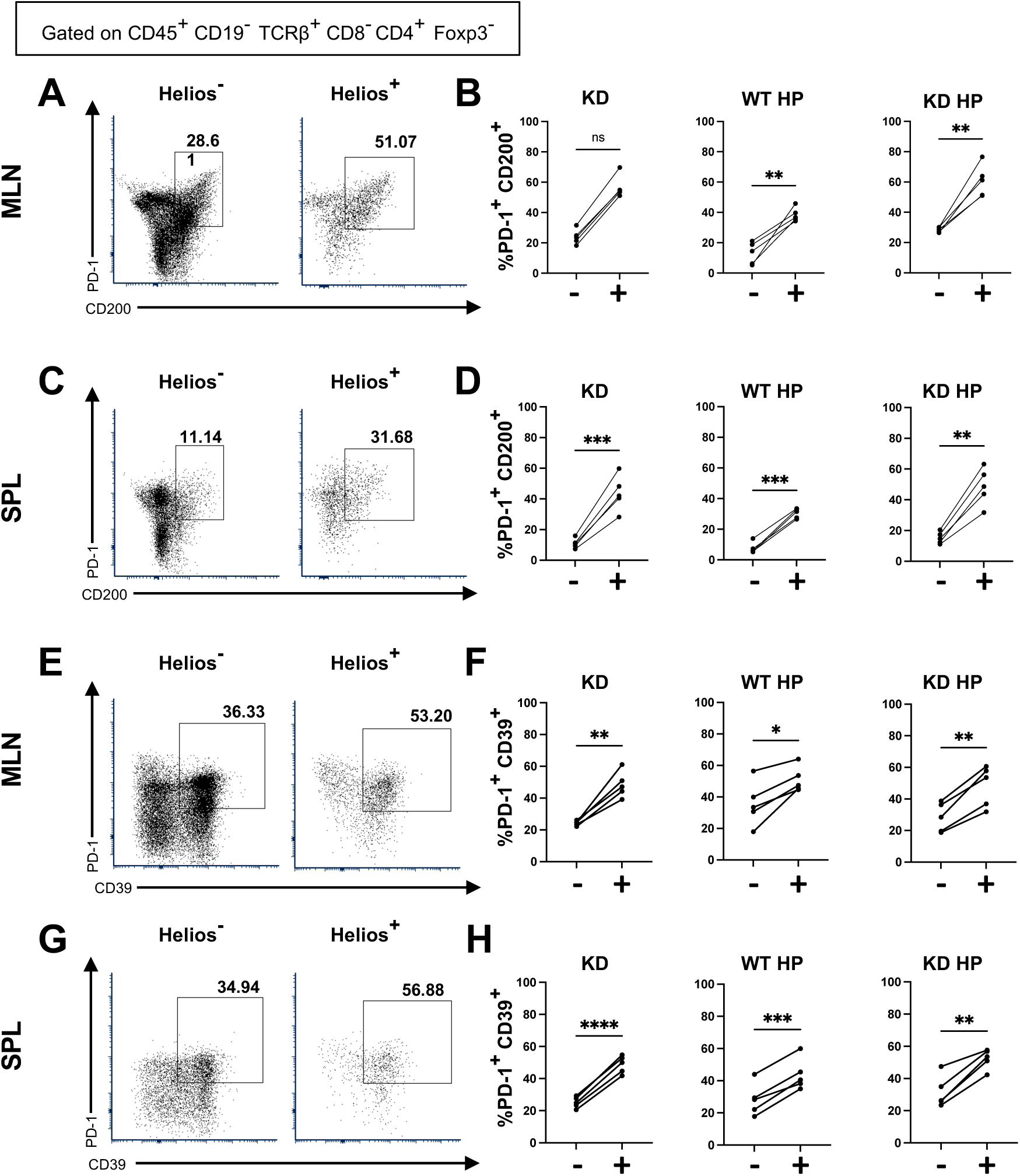
Systemic expression of PD-1, CD200 and CD39 on CD4^+^Helios^+^ cells. WT and CTLA4KD mice on the CB6F1 genetic background were administered HP by oral gavage at 1-3 months of age. Animals were sacrificed 6-8 months post HP infection. Single cell preparations were made from lymphoid organs and subsequently analyzed by flow cytometry. (A) Representative flow cytometry plots showing expression of PD-1 and CD200 in Helios^-^ cells (left) and Helios^+^ cells (right) from the MLN. (B) Quantification of PD-1 and CD200 co-expression in Helios^-^ (-) cells and Helios^+^ (+) cells from the MLN. (C) Representative flow cytometry plots showing expression of PD-1 and CD200 in Helios^-^ cells (left) and Helios^+^ cells (right) from the SPL. (D) Quantification of PD-1 and CD200 co-expression in Helios^-^ (-) cells and Helios^+^ (+) cells from the SPL. (E) Representative flow cytometry plots showing expression of PD-1 and CD39 in Helios^-^ cells (left) and Helios^+^ cells (right) from the MLN. (F) Quantification of PD-1 and CD39 co-expression in Helios^-^ (-) cells and Helios^+^ (+) cells from the MLN. (G) Representative flow cytometry plots showing expression of PD-1 and CD39 in Helios^-^ cells (left) and Helios^+^ cells (right) from the SPL. (H) Quantification of PD-1 and CD39 co-expression in Helios^-^ (-) cells and Helios^+^ (+) cells from the SPL. Each line represents one animal (*p ≤ 0.05, **p ≤ 0.01, ***p ≤ 0.001. ****p ≤ 0.0001)

**Supplemental Figure 6.**
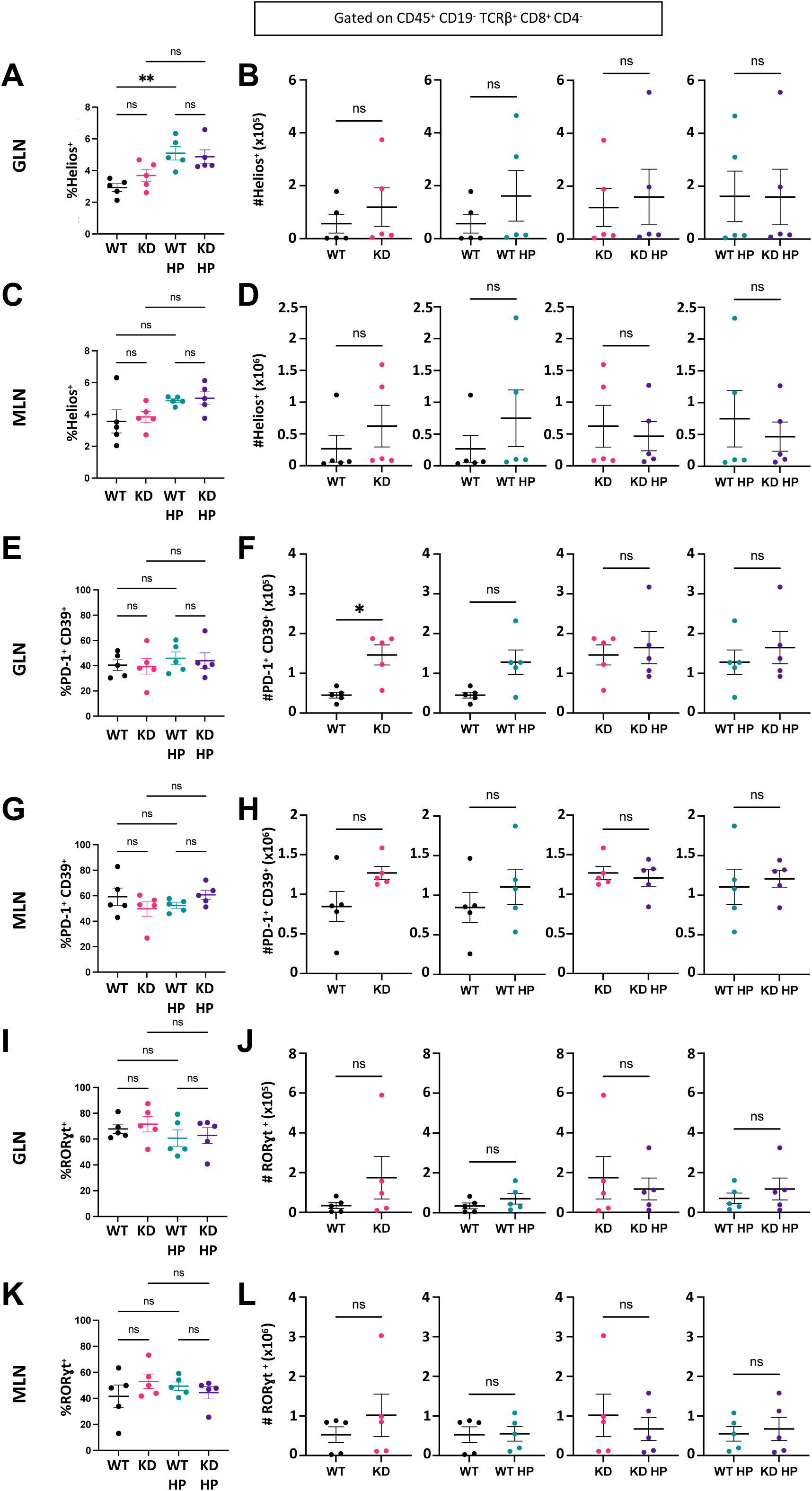
CD8 modulation by HP in the lymph nodes. WT and CTLA4KD mice on the B6 genetic background were administered HP by oral gavage at ∼4 weeks of age. Animals were sacrificed 2-3 months post HP infection. Single cell preparations were made from lymphoid organs and subsequently analyzed by flow cytometry. (A) Percentage of GLN CD8 T cells expressing Helios. (B) Number of Helios^+^ CD8 cells in the GLN. (C) Percentage of MLN CD8 T cells expressing Helios. (D) Number of CD8^+^Helios^+^ cells in the MLN. (E) Percentage of CD8 T cells co-expressing PD-1 and CD39 in the GLN. (F) Number of CD8 cell numbers co-expressing PD-1 and CD39 in the GLN. (G) Percentage of CD8 T cells co-expressing PD-1 and CD39 in the MLN. (H) Number of CD8 cell numbers co-expressing PD-1 and CD39 in the MLN. (I) Percentage of CD8 T cells expressing RORγt in the GLN. (J) Number of CD8 cell numbers expressing RORγt in the GLN. (K) Percentage of CD8 T cells expressing RORγt in the MLN. (L) Number of CD8 cell numbers expressing RORγt in the MLN. Each data point represents one animal (Mean± SEM, **p ≤ 0.01)

**Supplemental Figure 7.**
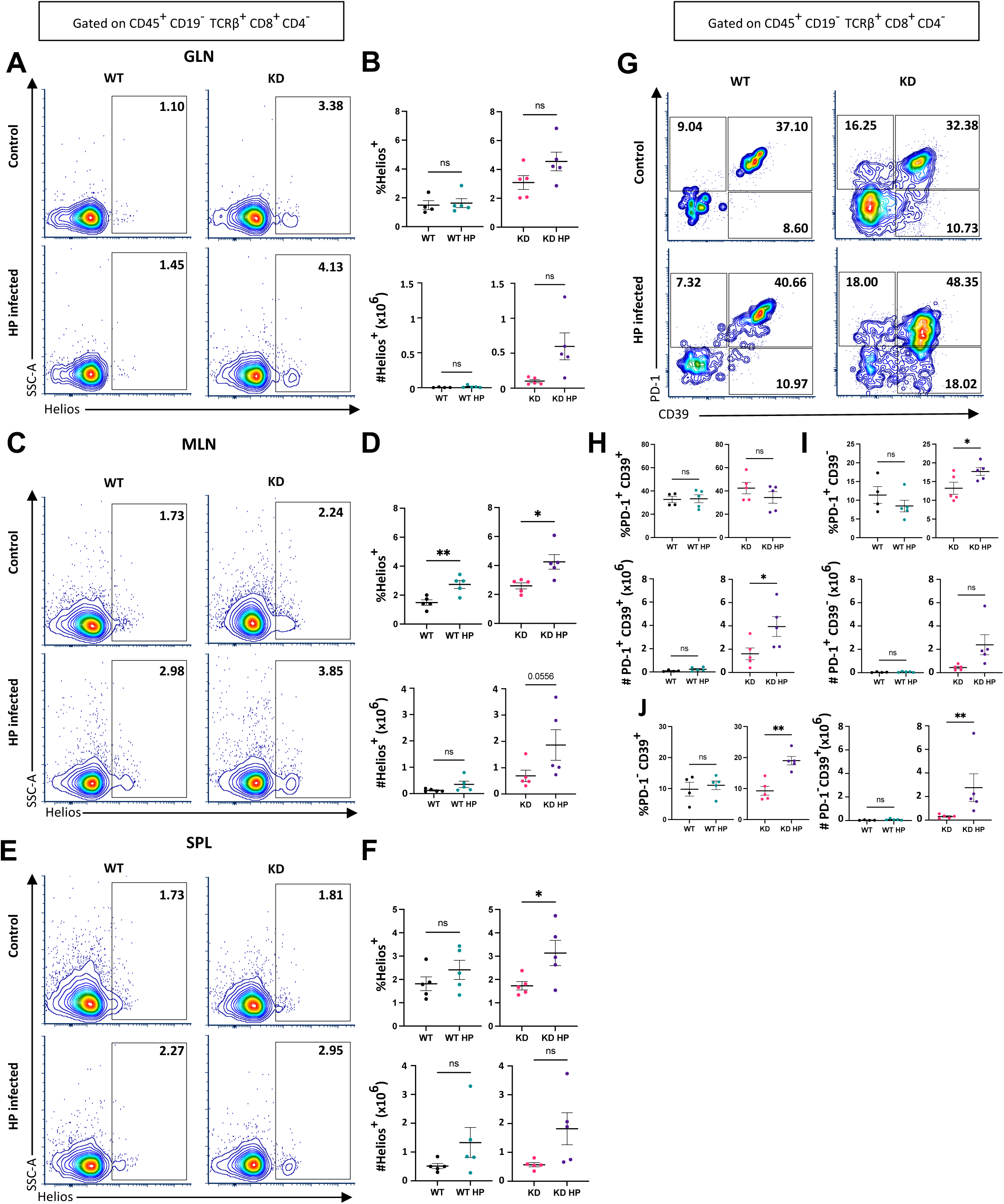
Systemic expression of Helios and local expression of PD-1 and CD39 in CD8 cells. WT and CTLA4KD mice on the CB6F1 genetic background were administered HP by oral gavage at 1-3 months of age. Animals were sacrificed 6-8 months post HP infection. Single cell preparations were made from lymphoid organs and subsequently analyzed by flow cytometry. (A) Representative flow cytometry plots displaying Helios and SSC-A in CD8^+^ T cells from the GLN. (B) Proportion of cells (top) and cell numbers (bottom) of CD8^+^ T cells from the GLN expressing Helios. (C) Representative flow cytometry plots displaying Helios and SSC-A in CD8^+^ T cells from the MLN. (D) Proportion of cells (top) and cell numbers (bottom) of CD8^+^ T cells from the MLN expressing Helios. (E) Representative flow cytometry plots displaying Helios and SSC-A in CD8^+^ T cells from the SPL. (F) Proportion of cells (top) and cell numbers (bottom) of CD8^+^ T cells from the SPL expressing Helios. (G) Representative flow cytometry plots displaying expression of PD-1 and CD39 on CD8^+^ T cells from the GLN. (H) Quantification of PD-1^+^CD39^+^ cell percentage (top) and cell numbers (bottom) from the GLN. (I) Quantification of PD-1^+^CD39^-^ cell percentage (top) and cell numbers (bottom) from the GLN. (J) Quantification of PD-1^-^CD39^+^ cell percentage (top) and cell numbers (bottom) from the GLN. Each data point represents one animal (Mean± SEM, *p ≤ 0.05, **p ≤ 0.01.)

## References

1. Thrift AP, Wenker TN, El-Serag HB. Global burden of gastric cancer: epidemiological trends, risk factors, screening and prevention. Nat Rev Clin Oncol 2023;20:338–349.

2. Ko KP. Risk Factors of Gastric Cancer and Lifestyle Modification for Prevention. J Gastric Cancer 2024;24:99–107.

3. Wroblewski LE, Peek RM, Jr., Wilson KT. Helicobacter pylori and gastric cancer: factors that modulate disease risk. Clin Microbiol Rev 2010;23:713–39.

4. Nguyen TH, Mallepally N, Hammad T, et al. Prevalence of Helicobacter pylori Positive Non-cardia Gastric Adenocarcinoma Is Low and Decreasing in a US Population. Dig Dis Sci 2020;65:2403–2411.

5. Sung H, Siegel RL, Rosenberg PS, et al. Emerging cancer trends among young adults in the USA: analysis of a population-based cancer registry. Lancet Public Health 2019;4:e137–e147.

6. Rawla P, Barsouk A. Epidemiology of gastric cancer: global trends, risk factors and prevention. Prz Gastroenterol 2019;14:26–38.

7. Song M, Latorre G, Ivanovic-Zuvic D, et al. Autoimmune Diseases and Gastric Cancer Risk: A Systematic Review and Meta-Analysis. Cancer Res Treat 2019;51:841–850.

8. Kehm RD, Yang W, Tehranifar P, et al. 40 Years of Change in Age- and Stage-Specific Cancer Incidence Rates in US Women and Men. JNCI Cancer Spectr 2019;3:pkz038.

9. Chambers CA, Kuhns MS, Egen JG, et al. CTLA-4-mediated inhibition in regulation of T cell responses: mechanisms and manipulation in tumor immunotherapy. Annu Rev Immunol 2001;19:565–94.

10. Schwab C, Gabrysch A, Olbrich P, et al. Phenotype, penetrance, and treatment of 133 cytotoxic T-lymphocyte antigen 4-insufficient subjects. J Allergy Clin Immunol 2018;142:1932–1946.

11. Egg D, Schwab C, Gabrysch A, et al. Increased Risk for Malignancies in 131 Affected CTLA4 Mutation Carriers. Front Immunol 2018;9:2012.

12. Hayakawa S, Okada S, Tsumura M, et al. A Patient with CTLA-4 Haploinsufficiency Presenting Gastric Cancer. J Clin Immunol 2016;36:28–32.

13. Miska J, Lui JB, Toomer KH, et al. Initiation of inflammatory tumorigenesis by CTLA4 insufficiency due to type 2 cytokines. J Exp Med 2018;215:841–858.

14. Merchant JL. Inflammation, atrophy, gastric cancer: connecting the molecular dots. Gastroenterology 2005;129:1079–82.

15. Hoft SG, Brennan M, Carrero JA, et al. Unveiling Cancer-Related Metaplastic Cells in Both Helicobacter pylori Infection and Autoimmune Gastritis. Gastroenterology 2025;168:53–67.

16. Fox JG, Wang TC, Rogers AB, et al. Host and microbial constituents influence Helicobacter pylori-induced cancer in a murine model of hypergastrinemia. Gastroenterology 2003;124:1879–90.

17. Honan AM, Jacobsen GE, Drum H, et al. Stromal-Like Cells Are Found in Peripheral Blood of Patients With Inflammatory Bowel Disease and Correlate With Immune Activation State. Clin Transl Gastroenterol 2024;15:e1.

18. Qiu P, Simonds EF, Bendall SC, et al. Extracting a cellular hierarchy from high-dimensional cytometry data with SPADE. Nat Biotechnol 2011;29:886–91.

19. Rogers AB. Histologic scoring of gastritis and gastric cancer in mouse models. Methods Mol Biol 2012;921:189–203.

20. Zhao H, Hu H, Chen B, et al. Overview on the Role of E-Cadherin in Gastric Cancer: Dysregulation and Clinical Implications. Front Mol Biosci 2021;8:689139.

21. Mori M, Shiraishi T, Tanaka S, et al. Lack of DMBT1 expression in oesophageal, gastric and colon cancers. Br J Cancer 1999;79:211–3.

22. Sautes-Fridman C, Petitprez F, Calderaro J, et al. Tertiary lymphoid structures in the era of cancer immunotherapy. Nat Rev Cancer 2019;19:307–325.

23. Sakaguchi S, Mikami N, Wing JB, et al. Regulatory T Cells and Human Disease. Annu Rev Immunol 2020;38:541–566.

24. Thornton AM, Korty PE, Tran DQ, et al. Expression of Helios, an Ikaros transcription factor family member, differentiates thymic-derived from peripherally induced Foxp3+ T regulatory cells. J Immunol 2010;184:3433–41.

25. Shafiei-Jahani P, Helou DG, Hurrell BP, et al. CD200-CD200R immune checkpoint engagement regulates ILC2 effector function and ameliorates lung inflammation in asthma. Nat Commun 2021;12:2526.

26. Mitchell P, Germain C, Fiori PL, et al. Chronic exposure to Helicobacter pylori impairs dendritic cell function and inhibits Th1 development. Infect Immun 2007;75:810–9.

27. Neyens D, Hirsch T, Abdel Aziz Issa Abdel Hadi A, et al. HELIOS-expressing human CD8 T cells exhibit limited effector functions. Front Immunol 2023;14:1308539.

28. Chellappa S, Hugenschmidt H, Hagness M, et al. CD8+ T Cells That Coexpress RORgammat and T-bet Are Functionally Impaired and Expand in Patients with Distal Bile Duct Cancer. J Immunol 2017;198:1729–1739.

29. Ihara T, Kushima R, Haruma K. Enhanced activity of autoimmune gastritis following Helicobacter pylori eradication therapy. Clin J Gastroenterol 2025;18:258–268.

30. Kim JH, Cheung DY. Must-Have Knowledge about the Helicobacter pylori-Negative Gastric Cancer. Gut Liver 2016;10:157–9.

31. Guilford P, Hopkins J, Harraway J, et al. E-cadherin germline mutations in familial gastric cancer. Nature 1998;392:402–5.

32. Murata-Kamiya N, Kurashima Y, Teishikata Y, et al. Helicobacter pylori CagA interacts with E-cadherin and deregulates the beta-catenin signal that promotes intestinal transdifferentiation in gastric epithelial cells. Oncogene 2007;26:4617–26.

33. Bruewer M, Luegering A, Kucharzik T, et al. Proinflammatory cytokines disrupt epithelial barrier function by apoptosis-independent mechanisms. J Immunol 2003;171:6164–72.

34. Saatian B, Rezaee F, Desando S, et al. Interleukin-4 and interleukin-13 cause barrier dysfunction in human airway epithelial cells. Tissue Barriers 2013;1:e24333.

35. Kang W, Nielsen O, Fenger C, et al. Induction of DMBT1 expression by reduced ERK activity during a gastric mucosa differentiation-like process and its association with human gastric cancer. Carcinogenesis 2005;26:1129–37.

36. Chen XY, Liu WZ, Shi Y, et al. Helicobacter pylori associated gastric diseases and lymphoid tissue hyperplasia in gastric antral mucosa. J Clin Pathol 2002;55:133–7.

37. Sato Y, Silina K, van den Broek M, et al. The roles of tertiary lymphoid structures in chronic diseases. Nat Rev Nephrol 2023;19:525–537.

38. Yang Y, Day J, Souza-Fonseca Guimaraes F, et al. Natural killer cells in inflammatory autoimmune diseases. Clin Transl Immunology 2021;10:e1250.

39. Henriques A, Teixeira L, Ines L, et al. NK cells dysfunction in systemic lupus erythematosus: relation to disease activity. Clin Rheumatol 2013;32:805–13.

40. Qin H, Lee IF, Panagiotopoulos C, et al. Natural killer cells from children with type 1 diabetes have defects in NKG2D-dependent function and signaling. Diabetes 2011;60:857–66.

41. Yun CH, Lundgren A, Azem J, et al. Natural killer cells and Helicobacter pylori infection: bacterial antigens and interleukin-12 act synergistically to induce gamma interferon production. Infect Immun 2005;73:1482–90.

42. Bamford KB, Fan X, Crowe SE, et al. Lymphocytes in the human gastric mucosa during Helicobacter pylori have a T helper cell 1 phenotype. Gastroenterology 1998;114:482–92.

43. Kao JY, Zhang M, Miller MJ, et al. Helicobacter pylori immune escape is mediated by dendritic cell-induced Treg skewing and Th17 suppression in mice. Gastroenterology 2010;138:1046–54.

44. Wang SK, Zhu HF, He BS, et al. CagA+ H pylori infection is associated with polarization of T helper cell immune responses in gastric carcinogenesis. World J Gastroenterol 2007;13:2923–31.

45. Ross EM, Bourges D, Hogan TV, et al. Helios defines T cells being driven to tolerance in the periphery and thymus. Eur J Immunol 2014;44:2048–58.

46. Koch MRA, Gong R, Friedrich V, et al. CagA-specific Gastric CD8(+) Tissue-Resident T Cells Control Helicobacter pylori During the Early Infection Phase. Gastroenterology 2023;164:550–566.

47. Li J, Zaslavsky M, Su Y, et al. KIR(+)CD8(+) T cells suppress pathogenic T cells and are active in autoimmune diseases and COVID-19. Science 2022;376:eabi9591.

